# Structural complementarity facilitates E7820-mediated degradation of RBM39 by DCAF15

**DOI:** 10.1101/738534

**Authors:** Tyler Faust, Hojong Yoon, Radosław P. Nowak, Katherine A. Donovan, Zhengnian Li, Quan Cai, Nicholas A. Eleuteri, Tinghu Zhang, Nathanael S. Gray, Eric S. Fischer

**Affiliations:** Department of Cancer Biology, Dana-Farber Cancer Institute, Boston, MA 02215, USA; Department of Biological Chemistry and Molecular Pharmacology, Harvard Medical School, Boston, MA 02115, USA

## Abstract

The investigational drugs E7820, indisulam and tasisulam (aryl-sulfonamides) promote the degradation of the splicing factor RBM39 in a proteasome-dependent mechanism. While the activity critically depends on the Cullin RING ligase substrate receptor DCAF15, the molecular details remain elusive. Here we present the cryo-EM structure of the DDB1-DCAF15-DDA1 core ligase complex bound to RBM39 and E7820 at 4.4 Å resolution, together with crystal structures of engineered subcomplexes. We show that DCAF15 adopts a novel fold stabilized by DDA1, and that extensive protein-protein contacts between the ligase and substrate mitigate low affinity interactions between aryl-sulfonamides and DCAF15. Our data demonstrates how aryl-sulfonamides *neo*-functionalize a shallow, non-conserved pocket on DCAF15 to selectively bind and degrade RBM39 and the closely related splicing factor RBM23 without the requirement for a high affinity ligand, which has broad implications for the *de novo* discovery of molecular glue degraders.

---

Pharmacologic intervention for many newly discovered disease targets — such as transcription factors, multi-protein complexes or scaffold proteins — is challenging because they lack an enzymatic function to facilitate the design of classical low molecular weight inhibitors. An alternative approach, small molecule-induced protein degradation, circumvents the need for an enzymatic function in the target protein^1^. The therapeutic potential of targeted protein degradation has been demonstrated by the success of thalidomide-related anti-cancer drugs (often referred to as immunomodulatory drugs, or IMiDs). IMiDs bind CRBN, the substrate receptor of the CUL4-RBX1-DDB1-CRBN (CRL4^CRBN^) E3 ubiquitin ligase^2–5^, and generate a novel binding surface to recruit and ubiquitinate *neo*-substrates^6–10^. Such molecular glues present an opportunity to target virtually any protein for degradation, even in the absence of a defined binding pocket. However, IMiDs have nanomolar affinity for CRBN, and the almost invariable conservation of the drug binding pocket and *neo*-substrate interaction surface suggests that IMiDs hijack an evolutionarily conserved mechanism, akin to what was found for the plant hormones auxin and jasmonate^11,12^. Whether molecular glue degraders critically depend on such high affinity interactions, and if these interactions can be achieved for ligases that have not evolved for ligand binding, is of critical importance for the further development of this new therapeutic modality.

Recently, the aryl-sulfonamides E7820, indisulam and tasisulam were shown to induce targeted degradation of the splicing factor RBM39 through recruitment of the E3 ubiquitin ligase CUL4-RBX1-DDB1-DCAF15 (CRL4^DCAF15^)^13,14^, which suggested a molecular glue mechanism. Indisulam was initially discovered in a phenotypic screen and found to be cytotoxic to specific cancer cell lines and in pre-clinical models^15^, while tasisulam and E7820 are derivatives around the sulfonamide core. E7820, indisulam and tasisulam were investigated in multiple phase I and II clinical trials involving advanced-stage solid tumors with a modest number of clinical responses, potentially due to an insufficient understanding of the mechanism of action and lack of informed patient stratification^14,16^. However, novel genetic dependencies in acute myeloid leukemia (AML) suggest a potential for clinical development^16^, and a recent phase II study encourages development with appropriate biomarkers^17^. Moreover, the aryl-sulfonamides appear to promote binding of DCAF15 to the RNA recognition motif (RRM) of RBM39, which suggests that derivatives of the aryl-sulfonamides may be used to target other RRM-containing proteins^9,10^. However, a detailed picture of the mechanism by which sulfonamides engage CRL4^DCAF15^ to promote turnover of the *neo*-substrate RBM39 is critically required to further leverage this new class of drugs for the targeting of RBM39, more generally of RRM containing proteins, and for the broad application of molecular glue degraders. We therefore set out to dissect the molecular basis of RBM39 recruitment to CRL4^DCAF15^.

## Results

### RBM39 recruitment to CRL4^DCAF15^ depends on sulfonamides

A recent study identified resistance mutations in cells treated with cytotoxic doses of indisulam that arise in the second RRM domain of RBM39 (RBM39_RRM2_)^13,14^. These mutations abrogate the interaction with CRL4^DCAF15^, which suggested that ligase binding is mediated by the RRM2 domain. To better characterize the interaction of RBM39 with DCAF15, we measured the affinity of recombinant DDB1-DCAF15 for RBM39_RRM2_ in the presence of E7820 using time-resolved fluorescence resonance energy transfer (TR-FRET). In the presence of E7820, indisulam or tasisulam at 50 µM, DDB1-DCAF15 and RBM39_RRM2_ associated with *K*_D_^app^ of 2.0 µM, 2.1 µM, and 3.5 µM, respectively (Fig. 1a, Supplementary Fig. 1a). In contrast, RBM39_RRM2_ did not show measurable affinity with DDB1-DCAF15, even at 10 µM, in the absence of compound (**Supplementary Fig. 1b**). E7820 interacts with DCAF15 (*K*_D_^app^ of 3.8 µM), but not with RBM39 (Fig. 1b, Supplementary Fig. 1c). Based on TR-FRET competition assays (**Supplementary Fig. 1d,e**), E7820 binds to DCAF15 with a *K*_*i*_ of 2.9 µM, while the *K*_*i*_ for indisulam and tasisulam is > 50 µM (Fig. 1c), which is analogous to the EC_50_ values when each compound is titrated into the RBM39_RRM2_ TR-FRET recruitment assay (**Supplementary Fig. 1f**). Notably, RBM39 was potently degraded in cells at 500 nM E7820 (**Supplementary Fig. 1g**), which contrasts the relatively weak affinity of E7820 for DCAF15.

**Fig. 1.**
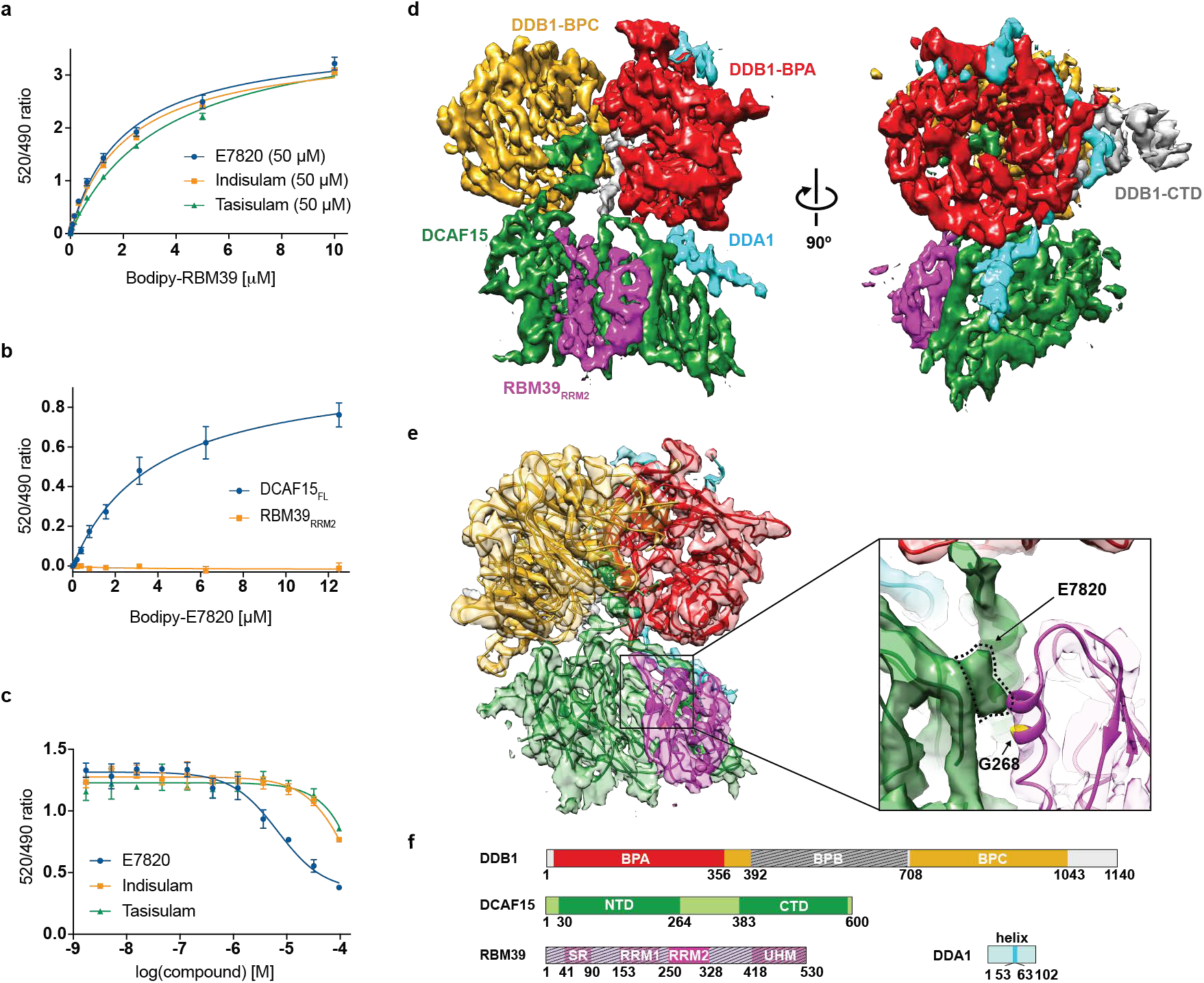
Cryo-EM structure of the DDB1∆B-DCAF15-DDA1 complex bound to E7820 and RBM39_RRM2_. **a**, TR-FRET. Titration of BodipyFL-RBM39_RRM2_ to DDB1∆B-DCAF15biotin in the presence of E7820 (**1**, *K*_D_^app^ = 2.0 µM), indisulam (**2**, *K*_D_^app^ = 2.1 µM), or tasisulam (**3**, *K*_D_^app^ = 3.5 µM) at 50 µM. **b**, TR-FRET. Titration of BodipyFL-E7820 (**4**) probe to DDB1∆B-DCAF15_biotin_ or RBM39_RRM2-biotin_. Compound binding is only observed for DDB1∆B-DCAF15_biotin_ (*K*_D_^app^ = 3.8 µM). **c**, Competitive titration of BodipyFL-E7820 (**4**) with aryl sulfonamides in TR-FRET assay. DDB1∆B-DCAF15_biotin_ is at 200 nM, BodipyFL-E7820 (**4**) is at 5 µM, and aryl-sulfonamides are at 0.002-100 µM. TR-FRET data in **a-c** are plotted as means ± s.d. from three independent replicates (*n* = 3). **d**, 4.4 Å cryo-EM map of the DDB1∆B-DCAF15-DDA1-E7820-RBM39_RRM2_ complex segmented to indicate DDA1 (cyan), DCAF15 (green), RBM39_RRM2_ (magenta), DDB1-BPC (orange), DDB1-BPA (red), and DDB1-CTD (grey). **e**, Cryo-EM map shown with the fitted and refined model. (Right), close-up of the region of the RBM39-DCAF15 interface, with the resistance mutation site G268V indicated in yellow and the putative E7820 density outlined in dotted lines. **f**, Domain representation of the proteins present in the complex. Regions omitted from the constructs are indicated by hatched lines.

### Cryo-EM structure of DCAF15 complex bound to RBM3_9RRM2_

All initial attempts to crystallize full-length human DCAF15 complexes were unsuccessful, so we focused our efforts on cryo-electron microscopy (cryo-EM). Initial class averages of DDB1-DCAF15-E7820-RBM39_RRM2_, indicated that DCAF15 and the BPB domain of DDB1 were flexible with respect to the core of DDB1 (**Supplementary Fig. 2a-d**). We therefore took advantage of a DDB1 construct lacking the BPB domain, DDB1∆B^18^, and chemical crosslinking (**Supplementary Fig. 2e**). DDB1∆B-DCAF15-DDA1-RBM39_RRM2_ were co-expressed in the presence of E7820, and after extensive optimization (see **Online Methods**), we collected a dataset that led to a 3D reconstruction of the 180 kDa complex at an overall resolution of ~ 4.4 Å (Fig. 1d-f, Supplementary Fig. 2e-h, and Supplementary Fig. 3).

DDB1∆B was readily placed into the density using the crystal structure (pdb: 5fqd, chain A) as a model, and a search using the balbes-molrep pipeline^19^ located the RRM domain corresponding to RBM39_RRM2_ (Fig. 1e) but did not identify homologous structures in the putative full-length DCAF15 density. The map allowed for segmentation of the density and unambiguous assignment of density to DCAF15 and DDA1 (Fig. 1d,e). While the resolution was not sufficient to build an atomic model (**Supplementary Fig. 3a**), we were able to build an approximate poly-alanine trace of DCAF15 and DDA1 using additional information from cross-linking mass spectrometry (**Supplementary Table 1**), mutations placed in putative helices (**Supplementary Fig. 4a**), and secondary structure prediction. RBM39_RRM2_ packs against an α-helix of DCAF15, and the Gly268 of RBM39, previously found to be a dominant position of indisulam resistance mutations^13,14^, packs against the DCAF15 helix and would not tolerate a sidechain-bearing residue (Fig. 1e). At the interface between RBM39_RRM2_ and DCAF15 was density that did not represent amino acid side chains and was tentatively assigned as E7820 (Fig. 1e). While the proximity of RBM39 residue Met265, which when mutated to leucine abrogates binding^14^, supports this assignment, the resolution of the cryo-EM map was insufficient for an unambiguous interpretation of ligand binding.

We therefore engineered a minimal complex suitable for crystallographic studies. Limited proteolysis experiments revealed that similarly sized fragments of DCAF15 were stably associated with DDB1 after gel filtration (**Supplementary Fig. 4b**). This result indicated both that DCAF15 contained an exposed, likely disordered, region available for proteolytic cleavage and that distinct segments of DCAF15 could independently bind DDB1. Disorder prediction further demonstrated a highly unstructured region of DCAF15 (**Supplementary Fig. 4c**), which led us to design constructs of the N-terminal (residues 30-264) and C-terminal (residues 383-600) fragments of human DCAF15 (DCAF15_split_). Co-expression of these fragments with DDB1∆B led to the formation of a soluble complex, that exhibited equivalent binding affinity for RBM39 to full-length human DCAF15 (**Supplementary Fig. 4d,e**).

### Crystal structure of DCAF15 complex bound to RBM39_RRM2_

Crystals were obtained for a DDB1∆B-DCAF15_split_-DDA1-E7820-RBM39_RRM2_ complex, and the structure was determined by molecular replacement with a final model refined to 2.9 Å resolution (Fig. 2a, **Supplementary Table 2**). To validate that the engineered DCAF15_split_ resembles the full-length DCAF15 structure, we docked the X-ray model into the cryo-EM map^20^ and found that the crystal structure accounts for all of the full-length DCAF15 density as well as density for E7820 (**Supplementary Fig. 3e,f**).

**Fig. 2.**
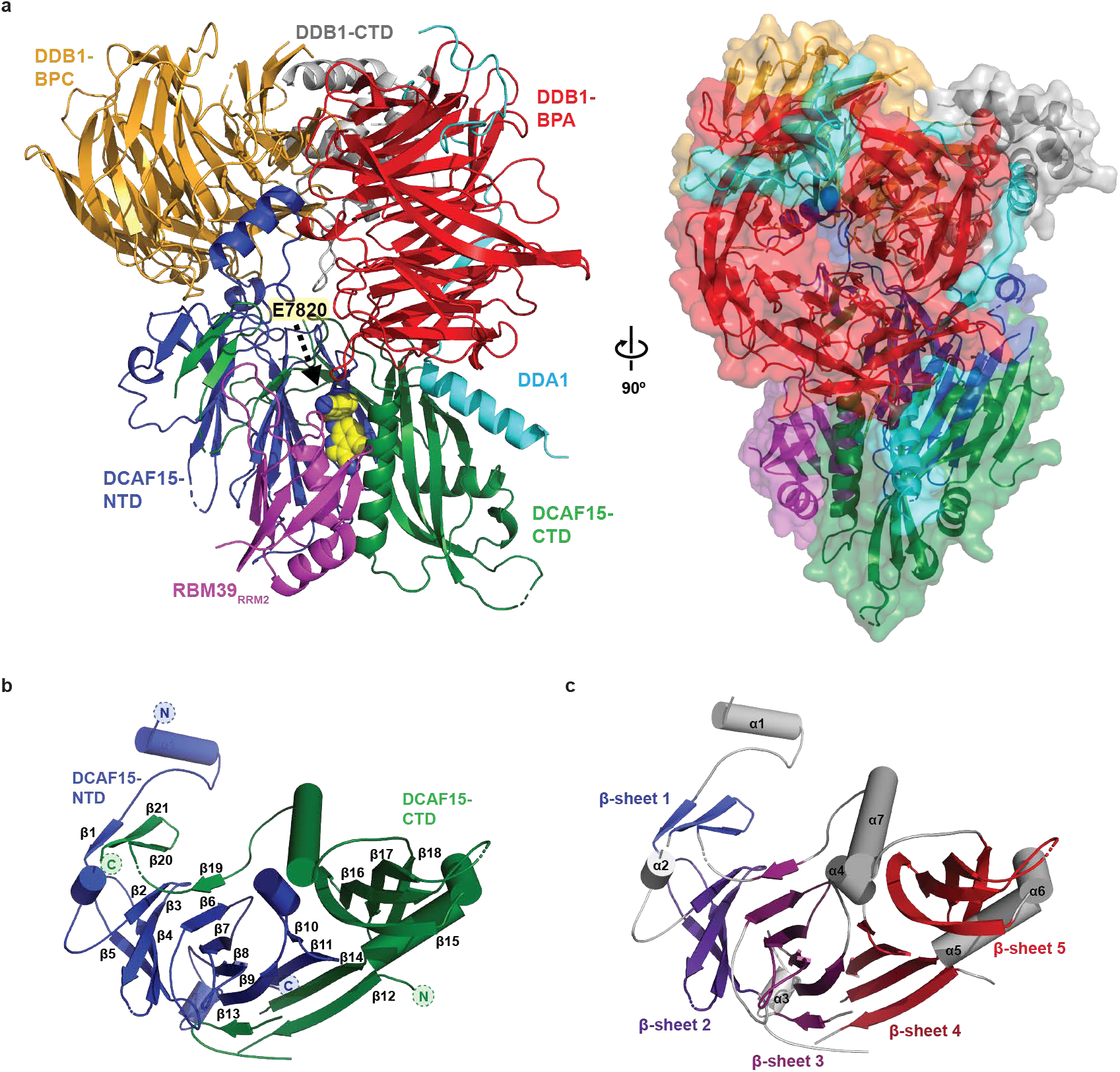
Crystal structure of the DDB1∆B-DCAF15_split_-DDA1-E7820-RBM39_RRM2_ complex. **a**, (Left) Cartoon representation of the DDB1∆B-DCAF15-DDA1-E7820-RBM39_RRM2_ complex. DDA1 (cyan), DCAF15-NTD (blue), DCAF15-CTD (green), RBM39_RRM2_ (magenta), DDB1-BPC (orange), DDB1-BPA (red), and DDB1-CTD (grey). E7820 is shown as spheres. (Right) A different view of the complex, shown in transparent surface representation. **b**, Cartoon representation of DCAF15 indicating secondary structure elements and colored in blue and green, for the DCAF15-NTD and DCAF15-CTD, respectively. DCAF15 alpha helices and beta strands are numbered from the N-to C-terminus, which are shown as colored circles for both the NTD and CTD of DCAF15. **c**, Cartoon view of DCAF15, highlighting the five stacked β**-**sheets. Helices from the NTD and CTD are colored in grey.

DCAF15_split_ consists of two predominantly β-sheet containing domains (Fig. 2b,c), the N-terminal domain (NTD, residues 30-264) and the C-terminal domain (CTD, residues 383-600). DCAF15 binds to DDB1 with a helix-loop-helix motif^21^, forming contacts with the two DDB1 β-propeller domains BPA and BPC and resembling the helix-loop-helix motif in CSA and DDB2 (**Supplementary Fig. 4a**, **Supplementary Fig. 5a,b**). DCAF15, unlike most other DDB1 and CUL4-associated factors (DCAFs), does not contain a canonical WD40 β-propeller fold and lacks homology to any other CRL substrate receptor^22^. Following the helix-loop-helix motif, the DCAF15 NTD and CTD are interwoven into five stacks of antiparallel β-sheets in an open solenoid arrangement, with β-sheets 1, 3, and 4 sharing strands from both the NTD and CTD. While β-sheets 2 and 3 have some resemblance to WD40 repeats, β-sheets 4 and 5 have unique features (Fig. 2b,c). Preceding β-sheet 4 is a short helix (α4) angled ~ 45° away from the sheet, before looping into β-strand 10 and 11. The terminal strands 12 and 14 of β-sheet 4 are contributed by the DCAF15 CTD, creating an extended interface between the two domains. β-sheet 5 is stabilized by two α-helices (DCAF15 α5 and α6), and α7 helix sits on the opposite side forming the major interactions with RBM39_RRM2_. The overall shape of DCAF15 is clamp-like and embraces RBM39_RRM2_ on the concave surface.

The small protein DDA1 is commonly associated with CRL4 complexes^23,24^, and knockout of DDA1 was found to reduce the indisulam-mediated degradation of RBM39^14^. In the crystal and cryo-EM structures, DDA1 binds to the top of the DDB1 BPA before running down the backside of the propeller (Fig. 1d, Fig. 3a). At the bottom of the DDB1 BPA, DDA1 intercalates a β-strand in the DDB1 propeller, using several highly conserved residues (Fig. 3b). Adjacent to this β-strand is an α-helix that buries multiple DDA1 hydrophobic residues (Leu55, Leu56, Leu59, and Trp63) in DCAF15 (Fig. 3b). Given that DDA1 is a core CRL4 component associating with many different substrate receptors^23,25^, the extent of the DCAF15 interactions are unexpected and suggest that the DDA1 helix represents a plastic binding module for other DCAFs. We measured the affinity of E7820 to recombinant DDB1-DCAF15 and DDB1-DCAF15-DDA1, as well as the ability of these complexes to bind to RBM39_RRM2_. While the affinity of E7820 to DCAF15 was not altered by the presence of DDA1, the apparent affinity to RBM39_RRM2_ was strengthened ~ 3-fold with an *K*_D_^app^ of 0.62 µM (Fig. 3c-e), which explains why genetic loss of DDA impairs induced RBM39 degradation^14^.

**Fig. 3.**
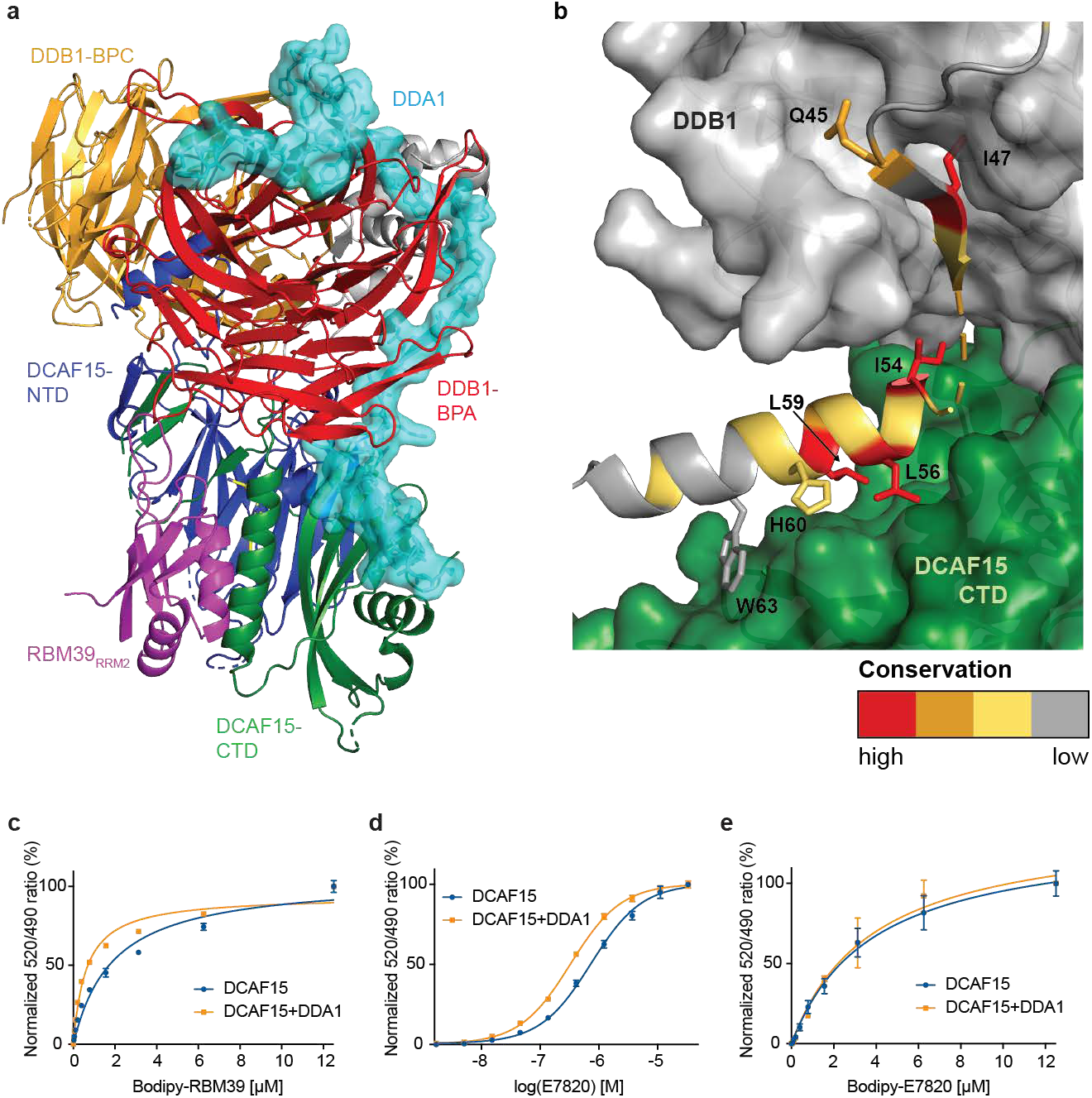
DDA1 stabilizes the CRL4^DCAF15^ complex and facilitates RBM39 recruitment. **a**, Cartoon representation of the DDB1∆B-DCAF15_split_-E7820-RBM39 complex with DDA1 highlighted as a cyan surface representation. DDA1 binds at the top of DDB1-BPA, winds down the back side of the propeller, and ends in a helix buried in DCAF15. **b**, DDB1 and DCAF15 are shown as a grey and green surface, respectively, and DDA1 is represented as a cartoon colored according to the conservation scores as calculated in ConSurf^36^. The top 3 bins of conservation in ConSurf (high conservation) are colored in red, orange, and yellow, respectively, while the bottom 6 bins (average and variable conservation, shown as “low”) are colored in gray to highlight the most conserved surfaces. **c**, TR-FRET. Titration of BodipyFL-RBM39_RRM2_ to DDB1∆B-DCAF15_biotin_ (*K*_D_^app^ = 1.9 µM) or DDB1∆B-DCAF15_biotin_-DDA1 (*K*_D_^app^ = 0.62 µM) in the presence of E7820 (50 µM), demonstrating enhanced recruitment of RBM39_RRM2_ to the DDA1-containing complex. **d**, TR-FRET. Titration of E7820 to DDB1∆B-DCAF15_biotin_ (EC_50_ = 0.74 µM) or DDB1∆B-DCAF15biotin-DDA1 and BodipyFL-RBM39_RRM2_ (EC_50_ = 0.33 µM). **e**, Titration of BodipyFL-E7820 to DDB1∆B-DCAF15_biotin_ (*K*_D_^app^ = 3.8 µM) or DDB1∆B-DCAF15_biotin_-DDA1 (*K*_D_^app^ = 3.8 µM). TR-FRET data in **c-e** are plotted as means ± s.d. from three independent replicates (*n* = 3).

### Aryl-sulfonamides interact primarily with DCAF15

E7820 binds in a shallow pocket at the interface between DCAF15-NTD and DCAF15-CTD situated in a weakly conserved surface groove proximal to DDB1 (Fig. 4, Supplementary Fig. 5c-e). While the placement of E7820 is firmly supported by the electron density (**Supplementary Fig. 6a,b**), we further validated the arrangement of the ligand through anomalous diffraction and a UV-crosslinking probe (**Supplementary Fig. 6c-h**). E7820 is sandwiched in a hydrophobic pocket between DCAF15 and RBM39_RRM2_, with the indole facing Met265 of RBM39. Notably, the RBM39 Met265Leu mutation was found to confer resistance to E7820-mediated degradation^14^, which is in accordance with the sulfur-π interaction observed in the structure. The two sulfonyl oxygens of E7820 form hydrogen bonds with the backbone amide nitrogens of DCAF15 Ala234 and Phe235, while the indole nitrogen and sulfonamide nitrogen form extensive water-mediated hydrogen bonds with the sidechain oxygens of RBM39 Thr262 and Asp264. Additional hydrogen bonds between the indole nitrogen and backbone carbonyl oxygen of DCAF15 Phe231, together form the core pharmacophore. The C4 methyl of E7820 forms hydrophobic interactions with Val477 and Val556 of DCAF15 (Fig. 4a,c), and swapping the methyl for a hydrogen, as in indisulam or desmethyl-E7820, results in a significant loss of DCAF15 binding (**Supplementary Fig. 6i**). The phenyl ring forms a T-shaped π–π interaction with DCAF15 Phe235 and otherwise is situated in a spacious pocket allowing for structural diversity as observed in indisulam and tasisulam.

**Fig. 4.**
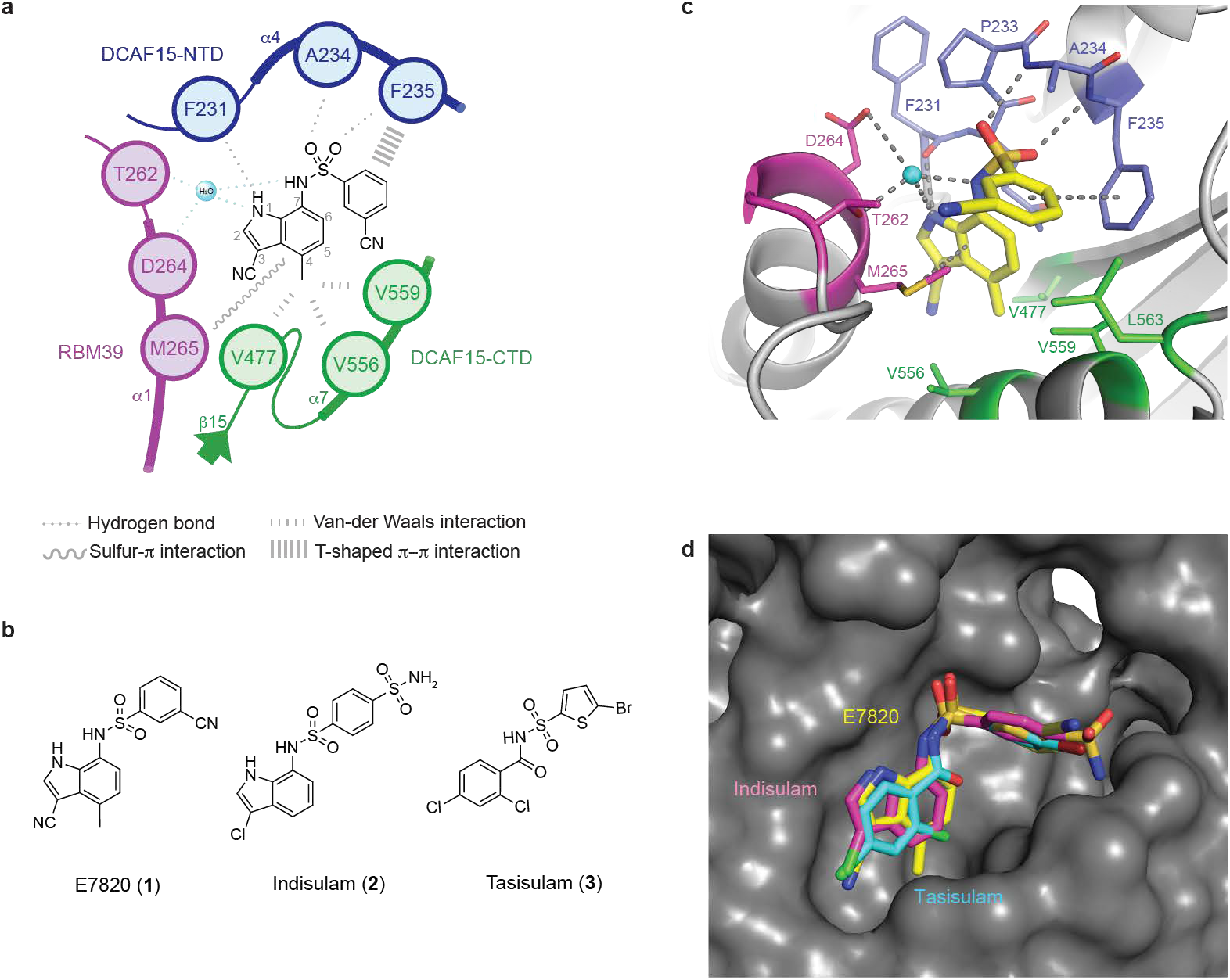
Aryl-sulfonamide binding to DCAF15. **a**, Sketch of E7820 and its interactions with DCAF15 and RBM39. Water-mediated hydrogen bonds are highlighted in cyan. **b**, Chemical structures of E7820 (**1**), indisulam (**2**), and tasisulam (**3**). **c**, E7820 interacts predominantly through the sulfonamide moiety and the indole moiety with residues in the DCAF15-NTD (blue). Additional hydrophobic interactions with the DCAF15-CTD (green), and sulfur-π interaction as well as water (cyan)-mediated hydrogen bonds with RBM39 (magenta) stabilize E7820 in a shallow pocket. **d**, Surface representation of DCAF15 is shown in grey and E7820, indisulam and tasisulam are shown as stick representation in yellow, magenta and cyan, respectively.

Finally, we obtained structures of the related but structurally distinct analogs indisulam and tasisulam to 2.9 Å resolution, respectively (Fig. 4d, Supplementary Fig. 6j,k). We find that indisulam and tasisulam bind DCAF15 in an overall configuration similar to E7820, maintaining the backbone hydrogen bonds from the sulfonyl groups to DCAF15 Ala234 and Phe235 and the water mediated hydrogen bonds. However, the methyl to hydrogen substitution at C4 in indisulam limits the hydrophobic interactions with DCAF15 Val477 and Val556, while tasisulam lacks the indole NH hydrogen bond to the backbone carbonyl of DCAF15 Phe231 (**Supplementary Fig. 6j,k**). These differences in indisulam and tasisulam help to explain their significant loss in affinity for DCAF15, while maintaining the ability to recruit RBM39 for degradation (Fig. 1a,c).

### DCAF15-RBM39 forms extensive protein-protein contacts

The weak affinity of aryl-sulfonamides for DCAF15 (**Supplementary Fig. 1c,f**) suggests that protein-protein contacts between DCAF15 and RBM39_RRM2_ stabilize the interaction. RBM39_RRM2_ presents itself as a canonical RRM fold, comprised of a four-stranded anti-parallel β-sheet (β1 - β4) stacked on two α-helices (α1 and α2) (Fig. 2a) and interacts with DCAF15 predominantly via the two α-helices. The RBM39_RRM2_ α1 helix docks into the surface groove on DCAF15 that also harbors the E7820 binding site and forms contacts with DCAF15 and E7820. The RBM39_RRM2_-DCAF15 interface comprises ~1,150 Å^2^ and spans the DCAF15 NTD and CTD (Fig. 5a). The binding groove is not conserved (**Supplementary Fig. 5e**) and is dominated by extensive hydrophobic interactions with the DCAF15 α7 helix in the CTD (Fig. 5b). As was observed in the cryo-EM structure (Fig. 1e), the tight packing of the interface would not allow a side chain-bearing residue at RBM39 Gly268, such that a Gly268Val mutation completely abrogates RMB39_RRM2_ recruitment to DCAF15 (**Supplementary Fig. 6l**). The interface includes four salt bridges between DCAF15 Arg574, Arg178, Arg160, and Asp174 and RBM39 Asp264, Glu271, and Arg275 respectively, and side chain hydrogen bonds between DCAF15 Ser546 and RBM39 Gln310, respectively (Fig. 5b). An additional indisulam resistance mutation in RBM39, Glu271Gln^14^, is likely explained by a loss in the salt bridge interaction with DCAF15 (**Supplementary Fig. 6m**). An extended network of backbone hydrogen bonds further stabilizes the DCAF15-RBM39 interface (**Supplementary Fig. 6n**).

**Fig. 5.**
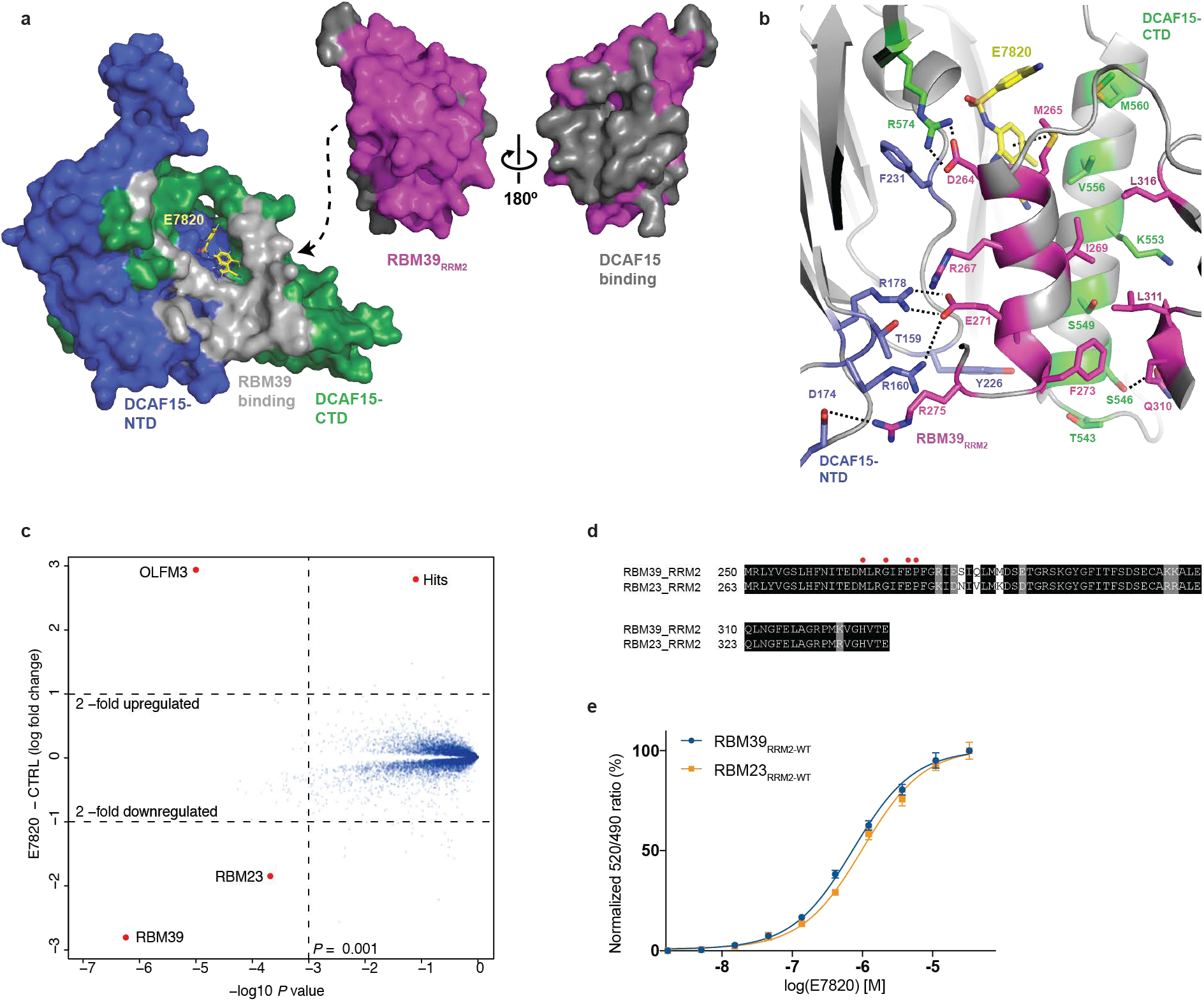
Inter-protein contacts between DCAF15 and RBM39. **a**, Surface representation of DCAF15 and RBM39_RRM2_ indicating the extensive interacting interface on DCAF15 and RBM39, shown in grey. E7820 is shown as a yellow stick representation. **b**, Side chain interactions between DCAF15, RBM39 and E7820. RBM39 buries a large hydrophobic surface on the DCAF15 α7 helix, in addition to four salt-bridges with DCAF15 on the opposing side of the binding interface. **c**, Scatter plot depicting identification of the novel E7820 substrate, RBM23, in Kelly cells. Kelly cells were treated with E7820 (10 µM) for 5 hours, and protein abundance was analyzed using TMT quantification mass spectrometry (two-sided moderated t-test as implemented in limma, *n = 3* for dmso, *n = 1* for E7820). **d**, Alignment of the second RRM domain from RBM39 and RBM23. Residues in black are completely conserved, gray shading represents similar substitutions, and white indicates no conservation. Red circles above the alignment indicate the positions of resistance mutations in RBM39 for indisulam-dependent toxicity. **e**, TR-FRET. Titration of E7820 to DDB1ΔB-DCAF15 in the presence of BodipyFL-RBM39_RRM2-WT_ (EC_50_ = 0.74 µM), BodipyFL-RBM23_RRM2-WT_ (EC_50_ = 1.0 µM). TR-FRET data in **e** are plotted as means ± s.d. from three independent replicates (*n* = 3).

### Aryl-sulfonamides selectively degrade of RBM39 and RBM23

As many RRM domains are structurally highly similar and since RBM39 interacts with DCAF15 predominantly through two conserved α-helices in its second RRM, we considered whether other RRM-containing proteins would be targeted by DCAF15 and E7820. To assess the degradome of E7820, we performed unbiased mass spectrometry-based proteomics experiments and found only RBM23 to be degraded in addition to RBM39 out of ~ 11,000 proteins detected (Fig. 5c). Sequence analysis revealed that the second RRM domain of RBM23 (RBM23_RRM2_) is nearly identical to RBM39_RRM2_, with 100% sequence identity across all key residues that form contacts with DCAF15 and E7820 (Fig. 5d). Consequently, we found comparable binding affinity for RBM23_RRM2_ to that observed for RBM39_RRM2_ (Fig. 5e). Cullin-RING ligases of the CRL4 family tolerate a diverse set of substrate receptors but typically present their substrates in a canonical position^21,26^. When superimposed with a Cullin-RING ligase complex (pdb: 4a0k), a model of the full CRL4^DCAF15^ ligase bound to RBM39 can be constructed. RBM39_RRM2_ is bound to a face of DCAF15 that is not directly opposed to RBX1 (Fig. 6a), however the N- and C-termini of RBM39 are positioned towards RBX1, and could tolerate additional domains at both positions. Furthermore, in contrast to CRBN, the ligand and substrate pocket of DCAF15 is not conserved (Fig. 6b), suggesting that the topological and evolutionary constraints on developing molecular glue degraders are rather flexible.

**Fig. 6.**
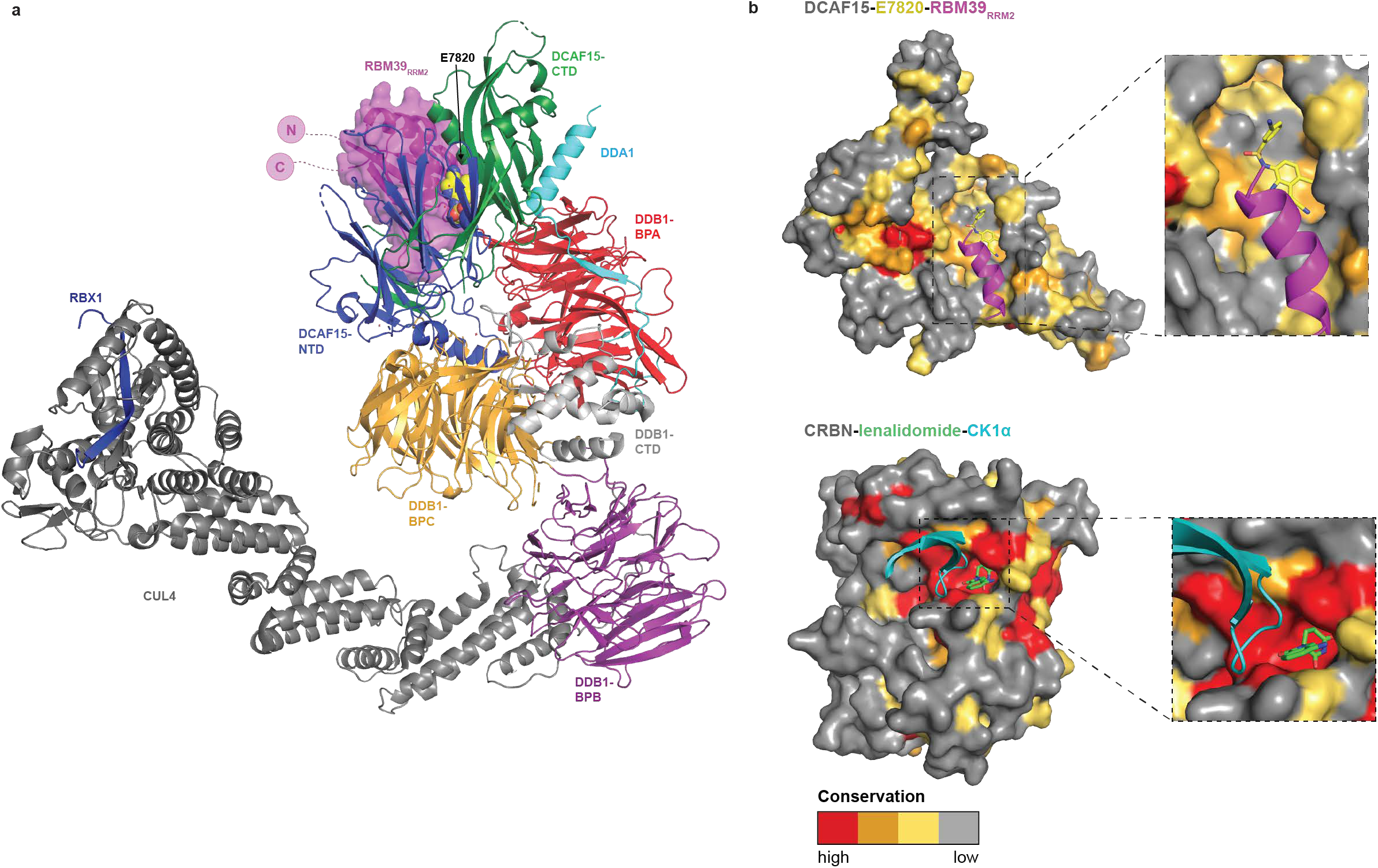
Topological and evolutionary constraints on E7820 activity. **a**, A model of the CRL4^DCAF15^ ligase bound to E7820 and RBM39_RRM2_. The N- and C-termini of RBM39_RRM2_ (pink circles) are positioned near RBX1 in the ligase, while RBM39_RRM2_ itself is bound on a non-proximal side face of DCAF15. The DCAF15_split_ crystal structure was superimposed onto the DDB1-DDB2-CUL4A-RBX1 crystal structure (pdb: 4a0k). **b**, Evolutionary conservation of DCAF15 (top) and CRBN (bottom). The substrate receptors are represented as a surface, colored according to the conservation scores as calculated in ConSurf with the top 3 bins of conservation colored in red, orange, and yellow, respectively, and the bottom 6 bins colored in gray to highlight the most conserved surfaces^36^. DCAF15 is shown bound to E7820 (yellow) and the α1 helix (residues 262-274) of RBM39_RRM2_ (magenta), while CRBN is shown bound to lenalidomide (green) and the β-hairpin loop (residues 29-49) of CK1α (cyan). Lenalidomide and CK1α both bind in a highly conserved pocket of CRBN.

## Discussion

Small molecules that recruit *neo*-substrates to a ubiquitin ligase have the potential to overcome the need for distinct binding pockets on target proteins. Our structural and pharmacological analyses have uncovered that aryl-sulfonamides represent a new type of molecular glue degrader, as they act as interface binders with weak receptor affinity to promote RBM39 degradation by CRL4^DCAF15^. We find that DCAF15 adopts a structure not commonly found in CRL substrate receptors and engages the second RRM domain of RBM39 with an extended, non-conserved surface area, which is in contrast to other small molecule-mediated CRL-substrate interactions such as auxin, jasmonate and IMiDs^10–12,18,27^. The total buried surface area of RBM39_RRM2_-DCAF15 is 1,150 Å^2^, which is larger than that of CRBN with *neo*-substrates (~600 Å^2^) and compensates for the weak affinity of these compounds for DCAF15 (*K*_*i*_ > 50 µM for indisulam and tasisulam). While IMiDs target a beta-hairpin loop primarily through backbone interactions between ligand, ligase and *neo*-substrate, the interactions between RBM39_RRM2_ and DCAF15 are more substantial and involve significant side-chain interactions that result in increased specificity compared to the relatively promiscuous zinc-finger recognition by the CRBN-IMiD complex^9,10^. The IMiD binding pocket is highly conserved, suggesting the existence of a natural ligand, while the pocket in DCAF15 is less conserved and implies that aryl-sulfonamides bind to an otherwise non-functional cavity. Together, the *neo*-functionalization of a relatively shallow, non-conserved pocket, and the weak affinity for DCAF15 suggests that molecular glues can be obtained for ligases that are not endogenously regulated by small molecules.

Induced protein degradation through PROTACs is another commonly used strategy for protein degradation and novel E3 ligase ligands are highly sought after for the development of such probes. The position of the binding pocket in DCAF15, which exposes the phenyl moiety to the solvent proximal to DDB1, together with the relatively weak affinity of aryl-sulfonamides for DCAF15, suggests that these ligands may not be ideally suited for the development of such degraders. While we demonstrate that a linker can be attached to the phenyl ring without significant loss of affinity, the exit vector is pointing towards DDB1 and away from RBX1, which will likely result in steric clashes with target proteins and sub-optimal positioning of recruited target proteins to be ubiquitinated.

Our work identifies DDA1 as an integral component of the CRL4^DCAF15^ ubiquitin ligase, and we demonstrate how DDA1 serves as an additional scaffolding subunit of a functional CRL4 complex. In the case of CRL4^DCAF15^, DDA1 binds to the top of the BPA subunit of DDB1 and forms extensive contacts with the backside of DDB1 before connecting to DCAF15 through an α-helix enforcing the overall structure of the complex. The presence of DDA1 causes increased E7820-dependent binding of DCAF15 to RBM39_RRM2_, which is in accordance with DDA1 knockout resulting in partial rescue of RBM39 degradation in cells^14^. It is conceivable that DDA1 will serve as a scaffolding protein in other CRL4 complexes, but more work is needed to understand the role of DDA1 in CRL4 regulation and its potential interplay with other CRL regulators such as the COP9 signalosome (CSN) and CAND1.

In summary, our work significantly expands our understanding of general principles of molecular glue degraders. We show how binders targeted to interfaces with significant sidechain interactions between receptor and substrate result in very selective agents. This is in contrast to IMiDs, which bind a conserved pocket in CRBN and only contribute minor backbone interactions to the interface, resulting in highly poly-targeted molecules^9,10^. While close analogs of E7820 are unlikely to target a broad set of RRM domains, the structural complementarity between DCAF15 and the canonical RRM fold suggests that novel chemical matter to target other RRM containing proteins to CRL4^DCAF15^ will likely be obtainable. Importantly, our structural characterization further supports the concept that compatible interfaces are more likely to occur between unrelated proteins than we may have anticipated. As such, prospective screens for molecular glues would ideally be preceded by selecting ligases with complementary interfaces for the intended target, similar to what we have previously utilized for PROTAC design^28^.

## Acknowledgements

We acknowledge Dr. Hyuk-Soo Seo for help with ITC experiments. Cryo-EM data was collected at the UMass cryo-EM facility, with help from Dr. KangKang Song and Dr. Chen Xu. Financial support for this work was provided by NIH grant NCI R01CA214608 (grant to E.S.F.) and F32 fellowship 1F32CA232772-01 (T.F.). E.S.F. is a Damon Runyon-Rachleff Innovator supported in part by the Damon Runyon Cancer Research Foundation (DRR-50–18). This work is based upon research conducted at the Northeastern Collaborative Access Team beamlines, which are funded by NIH NIGMS (P41 GM103403) and NIH-ORIP HEI grant (S10 RR029205). This research used resources of the Advanced Photon Source, a US Department of Energy (DOE) Office of Science User Facility operated by Argonne National Laboratory under Contract No. DE-AC02-06CH11357. This research was, in part, supported by the National Cancer Institute’s National Cryo-EM Facility at the Frederick National Laboratory for Cancer Research.

## Author contributions

T.F., H.Y., N.S.G., R.P.N., and E.S.F. initiated the project and designed experiments. T.F., H.Y. and R.P.N. conducted protein purification, T.F. performed crystallization and cryo-EM experiments. H.Y. developed and performed biochemical assays. R.P.N., T.F. and E.S.F. collected and processed X-ray diffraction data. N.A.E. and K.A.D. performed the mass spectrometry experiments. H.Y., Z.L., and Q.C. synthesized small molecules with input from T.Z. N.S.G. and E.S.F. supervised the project. T.F., H.Y., and E.S.F. wrote the manuscript, with input from all authors.

## Competing interests

E.S.F. is a founder and/or member of the scientific advisory board (SAB), and equity holder of C4 Therapeutics and Civetta Therapeutics and a consultant to Novartis, AbbVie and Pfizer. The Fischer lab receives research funding from Novartis, Deerfield and Astellas. N.S.G. is a founder, SAB member and equity holder in Gatekeeper, Syros, Petra, C4, B2S and Soltego. The Gray lab receives or has received research funding from Novartis, Takeda, Astellas, Taiho, Janssen, Kinogen, Voronoi, Her2llc, Deerfield and Sanofi. N.S.G., E.S.F, H.Y., Q.C., T.Z., T.F., R.P.N and K.A.D. are inventors on a patent application (PCT/US2018/065701 and PCT/US2019/014919), submitted by the Dana-Farber Cancer Institute.

## Online Methods

### Constructs and protein purification

The human genes for full-length DDB1, DDB1∆B (residues 396-705 replaced with GNGNSG linker), full-length DCAF15, DCAF15 NTD (30-264), DCAF15 CTD (383-600), full-length DDA1, RBM39_RRM2_ (245-332), and RBM23_RRM2_ (263-341) and the *Xenopus tropicalis* gene for full-length DCAF15 were cloned in pAC-derived vectors^29^. Baculovirus for protein expression (Invitrogen) was generated by transfection into *Spodoptera frugiperda* (Sf9) cells at a density of 0.9 × 10^6^ cells/mL grown in ESF 921 media (Expression Systems), followed by three rounds of infection in Sf9 cells to increase viral titer. Recombinant proteins were expressed as N-terminal His_6_, Strep II, Strep II Avi fusions in *Trichoplusia ni* High Five insect cells by infection with high titer baculovirus. Briefly, Hi Five cells grown in Sf-900 II SFM media (Gibco) at a density of 2.0 × 10^6^ cells/mL were infected with baculovirus at 1.5% (v/v). After 40 hours of expression at 27º C, Hi Five cells were pelleted for 10 minutes at 3,500 x g. For purification of StrepII or His_6_-tagged proteins, pelleted cells were resuspended in buffer containing 50 mM tris(hydroxymethyl) aminomethane hydrochloride (Tris–HCl) pH 8.0, 200 mM NaCl, 2 mM tris (2-carboxyethyl)phosphine (TCEP), 1 mM phenylmethylsulfonyl fluoride (PMSF), and 1× protease inhibitor cocktail (Sigma) and lysed by sonication. Media and purification buffers contained 10-20 µM E7820, as needed. Following ultracentrifugation, the soluble fraction was passed over the appropriate affinity resin of Strep-Tactin XT Superflow (IBA) or Ni Sepharose 6 Fast Flow affinity resin (GE Healthcare), eluted with wash buffer (50 mM Tris–HCl pH 8.0, 200 mM NaCl, 1 mM TCEP) supplemented with 50 mM d-Biotin (IBA) or 100 mM imidazole (Fisher Chemical), respectively. The affinity-purified DCAF15 complexes used for structure determination were next applied to an ion exchange column (Poros 50HQ) and eluted in 50 mM Tris-HCl pH 8.5, 2 mM TCEP, and 20 uM E7820 by a linear salt gradient (from 50-800 mM NaCl). Peak fractions of DCAF15 complex from ion exchange chromatography were then subjected to size-exclusion chromatography on a Superdex 200 10/300 in 50 mM 4-(2-hydroxyethyl)-1-piperazineethanesulfonic acid (HEPES) pH 7.4 or pH 8.0, 200 mM NaCl and 2 mM TCEP. Peak gel filtration fractions were pooled and concentrated and then either used directly in structural experiments or flash frozen in liquid nitrogen and stored at −80º C. Affinity-purified protein used in biochemical experiments was concentrated and subjected to size-exclusion chromatography as outlined above. The protein-containing fractions were concentrated using ultrafiltration (Millipore) and flash frozen in liquid nitrogen and stored at −80 °C.

### Limited proteolysis and gel filtration

The DDB1ΔB-*X.t.* DCAF15 complex was diluted to 20 µM in 25 mM HEPES pH 7.4, 200 mM NaCl, and 1 mM TCEP. *Xenopus tropicalis* DCAF15 is closely related to *Homo sapiens* DCAF15, with 66% sequence identity overall and 76% sequence identity in the structured NTD and CTD regions, and was examined in parallel in initial biochemical experiments. A 200 µM stock of chymotrypsin was diluted to 20 µM with 1 mM HCl and 2 mM CaCl_2_, which was then added to the DDB1ΔB-*X.t.* DCAF15 complex at a 400:1 ratio (50 nM chymotrypsin final concentration). The proteolysis reaction was carried out on ice for 45 minutes, centrifuged at 15,000 rpm at 4 °C, and injected onto an EnRich 650 column for gel filtration.

### Biotinylation of DCAF15 and RBM39

Purified Strep II Avi-tagged human DCAF15 variants or RBM39_RRM2_ were biotinylated *in vitro* at a concentration of 5-50 µM by incubation with final concentrations of 2.5 µM BirA enzyme and 0.2 mM D-Biotin in 50 mM HEPES pH 7.4, 200 mM NaCl, 10 mM MgCl_2_, 0.25 mM TCEP and 20 mM ATP. The reaction was incubated for 1 h at room temperature and stored overnight at 4 °C. Biotinylated proteins were purified by gel filtration chromatography and flash frozen in liquid nitrogen and stored at −80 °C.

### BodipyFL-labeling of RBM39 and RBM23

Purified human RBM39_RRM2_ or RBM23_RRM2_ was incubated with DTT (8 mM) at 4 °C for 1 h. DTT was removed using a S200 10/300 gel filtration column in a buffer containing 50 mM Tris pH 7.3 and 150 mM NaCl. BodipyFL-maleimide (Invitrogen) was dissolved in 100% DMSO and mixed with RBM39 or RBM23 to achieve 3-fold molar excess of BodipyFL-maleimide. Labelling was carried out at room temperature for 3 h and stored overnight at 4 °C. Labelled RBM39 or RBM23 was purified on a S200 10/300 gel filtration column in 50 mM Tris pH 7.5, 150 mM NaCl, 0.25 mM TCEP, concentrated by ultrafiltration (Milipore), flash frozen in liquid nitrogen and stored at −80 °C.

### Time-resolved fluorescence resonance energy transfer (TR-FRET)

Titrations of compounds to induce DCAF15-RBM39 or DCAF15-RBM23 complex were carried out by mixing 200 nM biotinylated Strep II Avi-tagged DCAF15, 200 nM BodipyFL-labeled RBM39 or RBM23 variants, and 2 nM terbium-coupled streptavidin (Invitrogen) in an assay buffer containing 50 mM Tris pH 8.0, 200 mM NaCl, 0.1% Pluronic F-68 solution (Sigma), and 0.5% BSA (w/v). Full-length human DCAF15 was used in all TR-FRET assays. After dispensing the assay mixture, increasing concentrations of compounds were dispensed in a 384-well microplate (Corning, 4514) using a D300e Digital Dispenser (HP) normalized to 2% DMSO. Before TR-FRET measurements were conducted, the reactions were incubated for 15 min at room temperature. After excitation of terbium fluorescence at 337 nm, emission at 490 nm (terbium) and 520 nm (BodipyFL) were recorded with a 70 μs delay over 600 μs to reduce background fluorescence, and the reaction was followed over 60 cycles of each data point using a PHERAstar FS microplate reader (BMG Labtech). The TR-FRET signal of each data point was extracted by calculating the 520/490 nm ratio. The half-maximal effective concentration EC_50_ values calculated using [Agonist] vs response (three parameters) equation in GraphPad Prism 7.

Titrations of BodipyFL-RBM39 were carried out by mixing 400 nM biotinylated Strep II Avi-tagged DCAF15 variants, 100 μM compounds or equivalent volume of DMSO, and 4 nM terbium-coupled streptavidin in the same assay buffer. After dispensing the assay mixture, increasing concentration of BodipyFL-RBM39 was added to the compound-bound DCAF15 in a 1:1 volume ratio and incubated for 15 min at room temperature. The 520/490 nm ratios were plotted to calculate the *K*_*d*_ values estimated using One site-Specific binding equation in GraphPad Prism 7.

Titrations of BodipyFL-E7820 (**4**) were carried out by mixing 200 nM biotinylated Strep II Avi-tagged DCAF15 variants or equivalent volume of the assay buffer, 2 nM terbium-coupled streptavidin in the same assay buffer. After dispensing the assay mixture, increasing concentration of BodipyFL-E7820 (**4**) was dispensed in the 384-well plate using D300e normalized to 2% DMSO, and then incubated for 15 min at room temperature. The 520/490 nm ratios from the sample with DCAF15 was subtracted by the ratios from the sample without DCAF15, and the subtracted values were plotted to calculate the *K*_*d*_ values estimated using One site-Specific binding equation in GraphPad Prism 7. All TR-FRET results are plotted as means ± s.d. from three independent replicates (*n* = 3) unless otherwise indicated.

### Crystallization

Frozen aliquots of the Strep II Avi-DCAF15_NTD_ (residues 30-264)-Strep II Avi-DCAF15_CTD_ (residues 383-600)-His_6_-DDB1ΔB-His_6_-DDA1-His_6_-RBM39_RRM2_ complex were thawed, centrifuged for 10 minutes at 15,000 rpm at 4 °C, and injected onto a Superdex 200 10/300 column equilibrated with 50 mM HEPES pH 8.0, 150 mM NaCl, 2 mM TCEP, and 20 μM E7820. All proteins used in crystallography are derived from human sequence. Peak fractions were pooled and concentrated at 4 °C to 56.8 μM (10 mg/mL). Concentrated protein was supplemented with 25 μM E7820, and crystallization plates were dispensed as sitting drop with the Formulatrix NT8 at room temperature. Crystals appeared within one day and continued growing until day 4 when concentrated protein was mixed 2:1 or 1:1 with reservoir containing 200 mM lithium citrate tribasic and 20% (w/v) PEG 3,350 in 96 well, 3 seat vapor diffusion Intelli-Plates (Art Robbins Instruments). For indisulam and tasisulam crystals, the same aliquots of Strep II Avi-DCAF15_NTD_ (residues 30-264)-Strep II Avi-DCAF15_CTD_ (residues 383-600)-His_6_-DDB1ΔB-His_6_-DDA1-His_6_-RBM39_RRM2_ complex bound to E7820 were thawed and diluted/concentrated two times with buffer containing 20 μM indisulam or 30 μM tasisulam, respectively. The first dilution was with 5-fold excess of gel filtration buffer containing the appropriate compound, and the second dilution was with 15-fold excess gel filtration buffer and compound. During the second dilution step, the protein complex was incubated on ice for 1 hour to allow complete exchange of the compound prior to concentration. After the second concentration step, protein complexes were injected onto a Superdex 200 10/300 column equilibrated with 50 mM HEPES pH 8.0, 150 mM NaCl, 2 mM TCEP and either 20 μM indisulam or 30 μM tasisulam. After gel filtration, purified protein was processed identically to E7820-bound complexes, as described above.

Crystals were cryo-protected in reservoir solution supplemented with 20% glycerol and flash frozen in liquid nitrogen. Diffraction data were collected at the APS Chicago (beamline 24-ID-C) with a Pilatus 6M-F detector at a temperature of 100 K, at wavelength of 0.9792 Å or 1.6531 Å. Data were indexed and integrated using XDS^30^ and scaled using AIMLESS supported by other programs of the CCP4 suite^31^. Data processing statistics, refinement statistics and model quality parameters are provided in **Supplementary Table 2**.

### Structure determination and model building

The DDB1ΔB-DCAF15_split_-DDA1-E7820-RBM39_RRM2_, DDB1ΔB-DCAF15_split_-DDA1-compound **5** (Iodide-E7820)-RBM39_RRM2_, DDB1ΔB-DCAF15_split_-DDA1-indisulam-RBM39_RRM2_, and DDB1ΔB-DCAF15_split_-DDA1-tasisulam-RBM39_RRM2_ complexes all crystallized in space group *P*2_1_2_1_2_1_with a single complex in the unit cell. PHASER32 was used to determine the structures by molecular replacement using a crystallographic model of DDB1∆B based on a crystal structure pdb: 5fqd. Diffraction data for complexes containing E7820-I or tasisulam were collected at 7500 eV and the MR-SAD pipeline as implemented in phaser^32^ used to obtain additional phase information, followed by density modification using parrot^31^. The initial model was iteratively improved with COOT^33^, using information from the density modified maps and sulfur anomalous difference peaks, and refined using PHENIX.REFINE^34^ and autoBUSTER^35^ with ligand restraints generated by Grade server (Global Phasing) or phenix.elbow^34^. Figures were generated with PyMOL (The PyMOL Molecular Graphics System, Version 2.3.0 Schrödinger, LLC) and model quality was assessed with MOLPROBITY. Interaction surfaces were determined with PISA, and conservation mapped using consurf^36^.

### Sample preparation and cryo-EM data collection

The DDB1-DCAF15-E7820-RBM39_RRM2_ complex was purified by gel filtration on a Superdex S200 10/300 column. A single peak fraction was collected and diluted to ~0.075 mg/mL. This diluted fraction was applied (4 μL) to a glow-discharged 1.2/1.3 Quantifoil copper 300 mesh grid, blotted for 3 seconds, and vitrified in liquid ethane with the Leica EM-GP blotting system. Micrographs were collected on a FEI Titan Krios at 300 kV, equipped with a K2 Summit camera and GIF energy filter. 1,457 micrographs were collected at the National Cryo-Electron Microscopy Facility (NCI) in super resolution mode at a pixel size of 0.532 Å. Each micrograph was recorded at a total dose of 40 e^−^/Å^2^ over 40 frames at a defocus range of 1.5-3.0 μm.

The DDB1ΔB-DCAF15-DDA1-E7820-RBM39_RRM2_ complex was purified by gel filtration, and peak fractions were pooled and concentrated for BS3 crosslinking. Briefly, 5 μM of complex was incubated with 60-fold molar excess of BS3 for 30 minutes at room temperature, quenched with 50 mM Tris-HCl pH 8.0, and re-injected on a Superdex 200 10/300. A peak fraction of crosslinked protein at ~0.048 mg/mL was applied (4 μL) to a glow-discharged 1.2/1.3 Quantifoil copper 300 mesh grid, blotted for 3 seconds, and vitrified in liquid ethane with the Lecia EM-GP blotting system. Data was collected from 2 grids over 4 imaging sessions on the same FEI Titan Krios at the UMass Cryo-EM facility, operating at 300 kV and equipped with a K2 Summit camera and GIF energy filter. The Volta phase plate (VPP) was used during all imaging sessions for this complex, and the position on the VPP was changed approximately every 400 micrographs. A total of 9,393 micrographs were collected in super resolution mode at a pixel size of 0.5294 Å. Each micrograph was recorded with a total dose of ~54 e^−^/Å^2^ over 35 or 40 frames, depending on the session. The defocus range was 0.2-2 μm across all micrographs.

### Image Processing

For the DDB1-DCAF15-E7820-RBM39_RRM2_ complex, all processing steps were performed in RELION 2. Movie frames were aligned and binned by a factor of 2 yielding a final pixel size of 1.064 Å and averaged with MotionCor2^37^, and CTF parameters were estimated with CTFFIND4^38^. A set of 1,000 particles were manually picked to generate 2D class averages for autopicking. Initial 2D classification was used to generate a starting set of 318,187 particles. From this set, two subsequent rounds of 3D classification with 7.5 degree angular sampling resulted in 68,324 particles for the final refinement, resulting in a reconstruction at 10 Å.

For the DDB1ΔB-DCAF15-DDA1-E7820-RBM39_RRM2_ complex, movie frames were aligned and binned by a factor of 2 yielding a final pixel size of 1.059 Å and averaged with MotionCor2^37^ and CTF parameters as well as the estimated phase shift were determined with CTFFIND4^38^. For the first three imaging sessions, ~5,000 particles were picked from each session to generate reference-free 2D class averages for automated picking in Relion. For the fourth session, crYOLO^39^ was used to pick particles with a model that was trained on the data. All subsequent processing steps for all sessions were performed with Relion 3.0^40^. Initial 2D classification was used to clean the data from each session independently, after which particles were pooled for further 3D classification. A round of 3D classification at 7.5 degree sampling was used to remove additional bad particles from the dataset, after which a set of 923,678 particles were used for CTF refinement and Bayesian polishing^40^. An initial round of CTF refinement on a consensus 3D refinement from all particles was performed to fit per-particle defocus. Thereafter, Bayesian polishing was performed independently on particles from each session. Particle images were then combined again, and it was found that an additional round of CTF refinement to estimate per-particle defocus led to an improved consensus 3D refinement. With the polished particles, one round of 3D classification with coarse (7.5 degree) angular sampling resulted in two main classes, one of which resulted in a reconstruction at 4.5 Å. The particles from this consensus refinement were further classified without image alignment, leading to a major class with 53% of the particles. 3D refinement of these particles improved the map quality, with a resolution of 4.5 Å. Finally, signal subtraction was performed on this consensus refinement with a soft subtraction mask around the DCAF15 CTD. An additional round of masked 3D classification without image alignment and a T value of 12, to account for the reduced signal in the particle box, again led to a dominant class with 56% of the particles. A final refinement with unsubtracted particles (75,529 particles in total) resulted in the final reconstruction at 4.4 Å. Local resolution was estimated using Relion.

### Cryo-EM model building

The refined and sharpened map from Relion^40^ was converted to structure factors using phenix map to structure factors^34^. DDB1∆B was placed using phenix dock in map, and the balbes-molrep pipeline^19^ used to place RBM39_RRM2_. The structure of the N-terminal region of DDA1 in complex with DDB1^24^ was used to trace DDA1. An approximate, partial poly-Ala model of DCAF15 was built in Coot^33^. First, well defined α-helices in the DCAF15 density were assigned based on secondary structure prediction, and mutations introduced to break helical fold or interactions (e.g. V43E and I45E in the putative helix-loop-helix motif anchoring DCAF15 to DDB1) and therefore further validate assignment. The remaining density was traced assisted by secondary structure predictions and distant constraints obtained through crosslinking mass spectrometry. Models were refined using phenix realspace refine^34^. To cross validate cryo-EM and X-ray structures, the final model obtained from the crystal structure was fitted into the cryo-EM volume using phenix dock in map, and subsequently realspace refined using phenix realspace refinement.

### Mutant DCAF15 pulldown

High five insect cells were infected with 1.5% (v/v) baculovirus expressing His_6_-DDB1ΔB, His_6_-RBM39_RRM2_, and wild type or mutant STREP II-DCAF15 full-length. After 40 hours, 1.5 mL of 50 mM Tris pH 8.0, 200 mM NaCl, 0.1% Triton X-100, 1 mM PMSF, 10 uM E7820, 2 mM TCEP and 1x protease cocktail (Sigma) was added to cell pellets and further lysed by sonication. Clarified lysates were then incubated with 50-100 μL of STREP-tactin XT superflow slurry (IBA), rocking at 4°C for one hour. Protein bound to STREP resin was washed 3x with 1 mL of lysis buffer and eluted with 2x packed bead volume of lysis buffer + 50 mM biotin. Eluted proteins were analyzed by SDS-PAGE.

### BS3, DSBU, DSSO cross-linking and MS

Recombinant DDB1-DCAF15-DDA1-E7820-RBM39_RRM2_ and DDB1ΔB-DCAF15-DDA1 were analyzed by the amine-reactive crosslinker DSSO and DSBU, while the DDB1-DCAF15-DDA1-E7820-RBM39_RRM2_ complex was also analyzed by BS3 crosslinking. For BS3 crosslinking, the protein complex was first injected onto a Superdex 200 10/300 and peak fractions were collected and concentrated to 1 mg/mL (4.6 μM) and 10 mM BS3 was added at 20, 40, 60, or 80x molar excess. Crosslinking reactions were incubated for 30 minutes at room temperature, followed by 5 minutes quench with 50 mM Tris-HCl pH 8.0. Similarly for DSSO and DSBU crosslinking, protein complexes were first injected onto a Superdex 200 10/300, peak fractions collected and concentrated to 10 μM. 50 mM of DSSO or DSBU was added at a 50, 100, or 200 molar excess. Crosslinking reactions were incubated for 30 minutes at room temperature, followed by 5 minutes quench with 20 mM Tris-HCl pH 8.0. All crosslinked samples were precipitated with tricholoracetic acid (TCA) following standard protocols^41^. Precipitated protein was then dissolved in 10 μL of 0.5 M Tris-HCl pH 8.6, 6 M guanidinium-hydrochloride and reduced, alkylated, and digested with either 200 ng trypsin or 600 ng chymotrypsin following standard protocols^42^. The digests were acidified with formic acid (ThermoFisher Scientific) and desalted using SOLAµ^TM^ SPE Plates (ThermoFisher Scientific).

Data were collected using an Orbitrap Fusion Lumos mass spectrometer (ThermoFisher Scientific, San Jose CA, USA) coupled with a Proxeon EASY-nLC 1200 LC pump (ThermoFisher Scientific). Peptides were separated on an EasySpray ES803 75 μm inner diameter microcapillary column (ThermoFisher Scientific). DSSO crosslinked peptides were separated using a 100 min gradient of 6–41% acetonitrile in 1.0% formic acid with a flow rate of 350 nL/min. The data were acquired using a mass range of *m/z* 375 – 1500, resolution 60,000, AGC target 4 × 10^5^, maximum injection time 50 ms, dynamic exclusion of 30 seconds for the peptide measurements in the Orbitrap. Data dependent MS2 spectra were acquired in the Orbitrap with a normalized collision energy (NCE) set at 25%, AGC target set to 5 × 10^4^ and a maximum injection time of 100 ms. For HCD-MS, MS2 fragment ions with a mass difference of 31.9721 Da (DSSO) or 26.0000 Da (DSBU) with 10-100% precursor intensity range were selected for fragmentation with HCD collision energy set to 30% and scans acquired in the Ion Trap with AGC target set to 2 × 10^4^, maximum injection time of 150 ms.

### Chemical crosslinking LC-MS data analysis

Proteome Discoverer 2.2 (ThermoFisher Scientific) with XLinkX version 2.2 was used for. RAW file processing and controlling peptide and protein level false discovery rates, assembling proteins from peptides, and protein quantification from peptides. MS/MS spectra were searched against a truncated (~200 proteins including the sequences for DCAF15, DDB1 and DDA1) Uniprot human database (September 2016) with both the forward and reverse sequences. Database search criteria are as follows: tryptic or chymotryptic with two missed cleavages, a precursor mass tolerance of 10 ppm, fragment ion mass tolerance of 0.6 Da, static alkylation of cysteine (57.0211 Da), variable oxidation of methionine (15.9951 Da), variable phosphorylation of serine, threonine and tyrosine (79.966 Da). DSSO crosslinked samples included the following variable modifications of lysines: DSSO (158.004 Da), amidated DSSO (142.050 Da) and hydrolysed DSSO (176.014 Da), and DSBU crosslinked samples included the following variable modifications of lysines: DSBU (196.085 Da), amidated DSBU (213.111 Da) and hydrolysed DSBU (214.095 Da).

### UV-crosslinking-coupled mass spectrometry

Purified DDB1∆B-DCAF15 full-length (3 µM) and His_6_-RBM39_RRM2_ (6 µM), and DMSO or E7820 (100 µM) were mixed and incubated for 15 min on ice. Compound **6** (Diazirine-E7820, 20 µM) or DMSO was added and incubated for 15 min on ice. The pre-mixed samples were irradiated with long-wave UV light for 15 min using a Spectrolinker UV Crosslinker (model XL1000, Spectronics Corp., Westbury, NY). The irradiated samples were processed as described above.

Data were collected using an Orbitrap Fusion Lumos mass spectrometer coupled with a Proxeon EASY-nLC 1200 LC pump. Peptides were separated on an EasySpray ES803 75 μm inner diameter microcapillary column. Peptides were separated using a 100 min gradient of 6– 38% acetonitrile in 1.0% formic acid with a flow rate of 350 nL/min. The data were acquired using a mass range of *m/z* 200 – 2000, resolution 120,000, AGC target 4 × 10^5^, maximum injection time 500 ms, dynamic exclusion of 60 seconds for the peptide measurements in the Orbitrap. Data dependent MS2 spectra were acquired in the ion trap with a normalized collision energy (NCE) set at 27%, AGC target set to 5 × 10^4^ and a maximum injection time of 100 ms.

Proteome Discoverer 2.2 was used to analyse the LC-MS data. MS/MS spectra were searched against a truncated (~200 proteins) Uniprot human database (September 2016) with both the forward and reverse sequences. Database search criteria are as follows: tryptic or chymotryptic with two missed cleavages, a precursor mass tolerance of 10 ppm, fragment ion mass tolerance of 0.02 Da, static alkylation of cysteine (57.0211 Da), variable oxidation of methionine (15.9951 Da), variable phosphorylation of serine, threonine and tyrosine (79.966 Da) and variable acetylation (42.011 Da) of the protein N-terminus and variable crosslinked compound **6** (possible adduct sizes: 422.141 Da or 83.049 Da) on all amino acids. Unique peptides were quantified in PD2.2 and the abundances of compound **6** modified peptides on DCAF15 for each of the treatments (DMSO, compound **6** (20 µM), and compound **6** (20 µM) + E7820 competition (100 µM)) were analysed for potential modification sites.

### TMT LC–MS3 mass spectrometry

Kelly cells were treated with DMSO vehicle (triplicate) or 10 µM of E7820 in singlicate for 5h. Treated Kelly cells were washed in PBS (Corning VWR, Radnor PA, USA) and collected at 3000 *g* centrifugation. Sample preparation and LC-MS analysis for whole proteome identification of novel E7820-dependent substrates was performed as described previously^9^.

### Data and materials availability

Structural coordinates for DDB1∆B-DDA1-DCAF15-E7820-RBM39, DDB1∆B-DDA1-DCAF15-tasisulam-RBM39, and DDB1∆B-DDA1-DCAF15-indisulam-RBM39 have been deposited in the Protein Data Bank under accession numbers 6Q0R, 6Q0V, and 6Q0W. The cryo-EM volume data are available at the EMDB, accession numbers: EMD-20554 and EMD-20553. Mass spectrometry raw data files have been deposited in PRIDE Archive under the accession numbers: PXD014536. Other data and materials are available from the authors upon reasonable request.

## References

1 Salami, J. & Crews, C. M. Waste disposal-An attractive strategy for cancer therapy. Science 355, 1163–1167, doi:10.1126/science.aam7340 (2017).

2 Chamberlain, P. P. et al. Structure of the human Cereblon-DDB1-lenalidomide complex reveals basis for responsiveness to thalidomide analogs. Nat Struct Mol Biol 21, 803–809 (2014).

3 Fischer, E. S., Park, E., Eck, M. J. & Thoma, N. H. SPLINTS: small-molecule protein ligand interface stabilizers. Curr Opin Struct Biol 37, 115–122, doi:10.1016/j.sbi.2016.01.004 (2016).

4 Fischer, E. S. et al. Structure of the DDB1-CRBN E3 ubiquitin ligase in complex with thalidomide. Nature 512, 49–53 (2014).

5 Ito, T. et al. Identification of a primary target of thalidomide teratogenicity. Science (New York, NY) 327, 1345–1350 (2010).

6 Lu, G. et al. The myeloma drug lenalidomide promotes the cereblon-dependent destruction of Ikaros proteins. Science 343, 305–309 (2014).

7 Kronke, J. et al. Lenalidomide causes selective degradation of IKZF1 and IKZF3 in multiple myeloma cells. Science 343, 301–305 (2014).

8 Gandhi, A. K. et al. Immunomodulatory agents lenalidomide and pomalidomide co-stimulate T cells by inducing degradation of T cell repressors Ikaros and Aiolos via modulation of the E3 ubiquitin ligase complex CRL4(CRBN.). British journal of haematology 164, 811–821 (2014).

9 Donovan, K. A. et al. Thalidomide promotes degradation of SALL4, a transcription factor implicated in Duane Radial Ray syndrome. Elife 7, doi:10.7554/eLife.38430 (2018).

10 Sievers, Q. L. et al. Defining the human C2H2 zinc finger degrome targeted by thalidomide analogs through CRBN. Science 362, doi:10.1126/science.aat0572 (2018).

11 Sheard, L. B. et al. Jasmonate perception by inositol-phosphate-potentiated COI1-JAZ co-receptor. Nature 468, 400–405, doi:10.1038/nature09430 (2010).

12 Tan, X. et al. Mechanism of auxin perception by the TIR1 ubiquitin ligase. Nature 446, 640–645, doi:10.1038/nature05731 (2007).

13 Uehara, T. et al. Selective degradation of splicing factor CAPERalpha by anticancer sulfonamides. Nat Chem Biol 13, 675–680, doi:10.1038/nchembio.2363 (2017).

14 Han, T. et al. Anticancer sulfonamides target splicing by inducing RBM39 degradation via recruitment to DCAF15. Science 356, doi:10.1126/science.aal3755 (2017).

15 Ozawa, Y. et al. E7070, a novel sulphonamide agent with potent antitumour activity in vitro and in vivo. Eur J Cancer 37, 2275–2282 (2001).

16 Wang, E. et al. Targeting an RNA-Binding Protein Network in Acute Myeloid Leukemia. Cancer Cell 35, 369–384 e367, doi:10.1016/j.ccell.2019.01.010 (2019).

17 Assi, R. et al. Final results of a phase 2, open-label study of indisulam, idarubicin, and cytarabine in patients with relapsed or refractory acute myeloid leukemia and high-risk myelodysplastic syndrome. Cancer 124, 2758–2765, doi:10.1002/cncr.31398 (2018).

18 Petzold, G., Fischer, E. S. & Thoma, N. H. Structural basis of lenalidomide-induced CK1alpha degradation by the CRL4 ubiquitin ligase. Nature 532, 127–130 (2016).

19 Brown, A. et al. Tools for macromolecular model building and refinement into electron cryo-microscopy reconstructions. Acta Crystallogr D Biol Crystallogr 71, 136–153, doi:10.1107/S1399004714021683 (2015).

20 Adams, P. D. et al. PHENIX: a comprehensive Python-based system for macromolecular structure solution. Acta Crystallographica Section D-Biological Crystallography 66, 213–221, doi:10.1107/S0907444909052925 (2010).

21 Fischer, E. S. et al. The molecular basis of CRL4DDB2/CSA ubiquitin ligase architecture, targeting, and activation. Cell 147, 1024–1039, doi:10.1016/j.cell.2011.10.035 (2011).

22 Zimmerman, E. S., Schulman, B. A. & Zheng, N. Structural assembly of cullin-RING ubiquitin ligase complexes. Current opinion in structural biology, doi:10.1016/j.sbi.2010.08.010 (2010).

23 Jin, J., Arias, E. E., Chen, J., Harper, J. W. & Walter, J. C. A family of diverse Cul4-Ddb1-interacting proteins includes Cdt2, which is required for S phase destruction of the replication factor Cdt1. Mol Cell 23, 709–721, doi:S1097-2765(06)00570-3[pii]10.1016/j.molcel.2006.08.010 (2006).

24 Shabek, N. et al. Structural insights into DDA1 function as a core component of the CRL4-DDB1 ubiquitin ligase. Cell Discov 4, 67, doi:10.1038/s41421-018-0064-8 (2018).

25 Olma, M. H. et al. An interaction network of the mammalian COP9 signalosome identifies Dda1 as a core subunit of multiple Cul4-based E3 ligases. Journal of Cell Science 122, 1035–1044, doi:10.1242/jcs.043539 (2009).

26 Cavadini, S. et al. Cullin-RING ubiquitin E3 ligase regulation by the COP9 signalosome. Nature 531, 598–603 (2016).

27 Matyskiela, M. E. et al. A novel cereblon modulator recruits GSPT1 to the CRL4(CRBN) ubiquitin ligase. Nature 535, 252–257 (2016).

28 Nowak, R. P. et al. Plasticity in binding confers selectivity in ligand-induced protein degradation. Nat Chem Biol, doi:10.1038/s41589-018-0055-y (2018).

## Methods-only References

29 Abdulrahman, W. et al. A set of baculovirus transfer vectors for screening of affinity tags and parallel expression strategies. Anal Biochem 385, 383–385 (2009).

30 Kabsch, W. Xds. Acta Crystallogr D Biol Crystallogr 66, 125–132, doi:10.1107/S0907444909047337 (2010).

31 Winn, M. D. et al. Overview of the CCP4 suite and current developments. Acta crystallographica. Section D, Biological crystallography 67, 235–242, doi:10.1107/S0907444910045749 (2011).

32 Mccoy, A. et al. Phaser crystallographic software. Journal of Applied Crystallography 40, 658–674 (2007).

33 Emsley, P. & Cowtan, K. Coot: model-building tools for molecular graphics. Acta Crystallographica Section D-Biological Crystallography 60, 2126–2132 (2004).

34 Afonine, P. V. et al. Towards automated crystallographic structure refinement with phenix.refine. Acta Crystallographica Section D 68, 352–367, doi:doi: 10.1107/S0907444912001308 (2012).

35 BUSTER version 2.10.2 v. 2.10.2 (Global Phasing Ltd., Cambridge, United Kingdom, 2011).

36 Landau, M. et al. ConSurf 2005: the projection of evolutionary conservation scores of residues on protein structures. Nucleic acids research 33, W299–302, doi:10.1093/nar/gki370 (2005).

37 Zheng, S. Q. et al. MotionCor2: anisotropic correction of beam-induced motion for improved cryo-electron microscopy. Nat Methods 14, 331–332, doi:10.1038/nmeth.4193 (2017).

38 Rohou, A. & Grigorieff, N. CTFFIND4: Fast and accurate defocus estimation from electron micrographs. J Struct Biol 192, 216–221, doi:10.1016/j.jsb.2015.08.008 (2015).

39 Moriya, T. et al. High-resolution Single Particle Analysis from Electron Cryo-microscopy Images Using SPHIRE. J Vis Exp, doi:10.3791/55448 (2017).

40 Zivanov, J. et al. New tools for automated high-resolution cryo-EM structure determination in RELION-3. Elife 7, doi:10.7554/eLife.42166 (2018).

41 Link, A. J. & LaBaer, J. Trichloroacetic acid (TCA) precipitation of proteins. Cold Spring Harb Protoc 2011, 993–994, doi:10.1101/pdb.prot5651 (2011).

42 Gundry, R. L. et al. Preparation of proteins and peptides for mass spectrometry analysis in a bottom-up proteomics workflow. Curr Protoc Mol Biol Chapter 10, Unit10 25, doi:10.1002/0471142727.mb1025s88 (2009).

