## Supplementary Information for "Structural complementarity facilitates E7820-mediated degradation of RBM39 by DCAF15"

| Protein 1 | Protein 2 | Distance (Å) | observ ations | apo | +RBM39 | Cross-linker |
|---|---|---|---|---|---|---|
| DCAF15-K85 | DDB1-K1081 | 14.1 | 3 | yes | | DSBU, DSSO |
| DDA1-K13 | DDB1-K335 | 14.1 | 2 | yes | | DSBU |
| DDA1-K66 | DDB1-K204 | 20 | 1 | yes | | DSBU |
| DCAF15-K56 | DCAF15-K85 | 20.2 | 1 | yes | | DSSO |
| DCAF15-K56 | DCAF15-K587 | 19.8 | 2 | yes | | DSBU |
| DCAF15-K511 | DCAF15-K540 | 6.7 | 3 | yes | | DSSO |
| DCAF15-K582 | DCAF15-K587 | 12.3 | 1 | yes | | DSSO |
| DDA1-K65 | DDA1-K70 | 15.3 | 2 | | yes | BS3 |
| DDB1-K11 | DDB1-K35 | 10 | 2 | yes | | DSBU, DSSO |
| DDB1-K35 | DDB1-K857 | 45.8 | 1 | yes | | DSBU |
| DDB1-K53 | DDB1-K1104 | 13.1 | 5 | yes | | DSBU, DSSO |
| DDB1-K70 | DDB1-K150 | 20.7 | 3 | yes | | DSBU, DSSO |
| DDB1-K70 | DDB1-K200 | 19.5 | 2 | yes | | DSBU |
| DDB1-K150 | DDB1-K200 | 13.5 | 2 | yes | | DSBU, DSSO |
| DDB1-K153 | DDB1-K200 | 6.9 | 1 | yes | | DSSO |
| DDB1-K191 | DDB1-K204 | 7.7 | 4 | yes | | DSBU, DSSO |
| DDB1-K244 | DDB1-K298 | 10 | 4 | yes | | DSBU, DSSO |
| DDB1-K769 | DDB1-K857 | 21.2 | 2 | yes | | DSBU, DSSO |
| DDB1-K769 | DDB1-K864 | 10.9 | 3 | yes | | DSBU, DSSO |
| DDB1-K769 | DDB1-K867 | 14.1 | 1 | yes | | DSSO |
| DDB1-K769 | DDB1-K897 | 18.5 | 1 | yes | | DSSO |
| DDB1-K823 | DDB1-K897 | 15.4 | 6 | yes | | DSBU, DSSO |
| DDB1-K857 | DDB1-K897 | 16.1 | 1 | yes | | DSBU |
| DDB1-K867 | DDB1-K897 | 25 | 1 | yes | | DSBU |
| DDB1-K917 | DDB1-K979 | 11 | 6 | yes | | DSBU, DSSO |
| DDB1-K936 | DDB1-K979 | 10.1 | 6 | yes | | DSBU, DSSO |
| DCAF15-K26 | RBM39-K291 | n/a | 2 | | yes | BS3 |
| DCAF15-K321 | RBM39-K178 | n/a | 1 | | yes | BS3 |
| DCAF15-K121 | DDB1-K628 | n/a | 1 | | yes | BS3 |
| DCAF15-K321 | DDB1-K864 | n/a | 1 | | yes | DSSO |
| DCAF15-K332 | DDB1-K287 | n/a | 1 | | yes | DSBU |
| DCAF15-K332 | DDA1-K65 | n/a | 2 | | yes | BS3 |
| DDA1-K26 | DDB1-K35 | n/a | 3 | yes | | DSBU |
| DDA1-K26 | DDB1-K53 | n/a | 2 | yes | | DSSO |
| DDA1-K89 | DDB1-K204 | n/a | 1 | yes | | DSSO |
| DCAF15-K6 | DCAF15-K26 | n/a | 9 | yes | yes | BS3, DSBU, DSSO |
| DCAF15-K26 | DCAF15-K38 | n/a | 3 | yes | yes | BS3, DSSO |
| DCAF15-K26 | DCAF15-K56 | n/a | 2 | | yes | BS3 |
| DCAF15-K319 | DCAF15-K332 | n/a | 5 | yes | yes | BS3, DSSO |
| DCAF15-K319 | DCAF15-K335 | n/a | 3 | yes | yes | DSSO |
| DCAF15-K321 | DCAF15-K332 | n/a | 11 | | yes | BS3, DSBU, DSSO |
| DCAF15-K321 | DCAF15-K335 | n/a | 10 | yes | yes | BS3, DSBU, DSSO |

| | | | | | | |
|---|---|---|---|---|---|---|
| DCAF15-K332 | DCAF15-K412 | n/a | 1 | | yes | BS3 |
| DCAF15-K335 | DCAF15-K511 | n/a | 1 | yes | | DSBU |
| DDA1-K26 | DDA1-K51 | n/a | 3 | | yes | BS3, DSBU |
| DDA1-K65 | DDA1-K71 | n/a | 1 | | yes | DSBU |
| DDA1-K65 | DDA1-K89 | n/a | 1 | yes | | DSSO |

**Supplementary Table 1. Table of lysine pairs identified by protein cross-linking.** Distances were measured in PyMOL using the crystal structure, except in cases where one or both lysines were absent from the crystal structure. Observations indicate the number of individual experiments where each cross-link was identified. The DCAF15-DDB1ΔB-DDA1 (apo) and DCAF15-DDB1-DDA1-E7820-RBM39$_{RRM2}$ (+RBM39) complexes were both analyzed by cross-linking mass spec, and the identification of cross-links in either complex is indicated. Also listed are the crosslinkers (BS3, DSSO, or DSBU) that resulted in each crosslink pair.

| | X102 E7820 | X180 indisulam | X198 tasisulam |
|---|---|---|---|
| **<u>Data collection</u>** | | | |
| Space group | $P2_12_12_1$ | $P2_12_12_1$ | $P2_12_12_1$ |
| Cell dimensions | | | |
| *a*, *b*, *c* (Å) | 81.1, 93.6, 258.43 | 93.97, 81.80, 260.98 | 81.77, 94.61, 260.40 |
| α, β, γ (°) | 90.0, 90.0, 90.0 | 90.0, 90.0, 90.0 | 90.0, 90.0, 90.0 |
| Resolution (Å) | 45.3–2.9 (3.0–2.9)* | 45.6–2.9 (3.0–2.9)* | 46.5–2.9 (3.0–2.9)* |
| $R_{sym}$ or $R_{merge}$ | 0.03 (0.80) | 0.17 (2.32) | 0.03 (0.72) |
| $I / \sigma I$ | 13.21 (0.79) | 8.19 (0.71) | 15.97 (0.78) |
| *CC 1/2* | 1.00 (0.69) | 0.99 (0.60) | 0.99 (0.82) |
| Completeness (%) | 99.5 (99.6) | 99.6 (98.6) | 99.3 (97.2) |
| Redundancy | 2.0 (2.0) | 6.7 (6.9) | 2.0 (2.0) |
| **<u>Refinement</u>** | | | |
| Resolution (Å) | 45.3–2.9 | 45.6–2.9 | 46.5–2.9 |
| No. reflections | 44327 | 45335 | 45140 |
| $R_{work}$ / $R_{free}$ | 21.5 / 25.8 | 21.2 / 23.7 | 20.3 / 25.1 |
| No. atoms | | | |
| Protein | 10180 | 10314 | 10213 |
| Ligand/ion | 25 | 25 | 21 |
| Water | 11 | 3 | |
| *B*-factors | | | |
| Protein | 131.5 | 117.1 | 116.9 |
| Ligand/ion | 99.2 | 111.7 | 95.8 |

| | | | |
|---|---|---|---|
| Water | 81.6 | 79.1 | |
| **R.m.s. deviations** | | | |
| Bond lengths (Å) | 0.014 | 0.014 | 0.014 |
| Bond angles (°) | 1.92 | 1.86 | 1.93 |

* Values in parentheses are for highest-resolution shell.

**One crystal was used for each structure.

**Supplementary Table 2.** Data collection and refinement statistics.

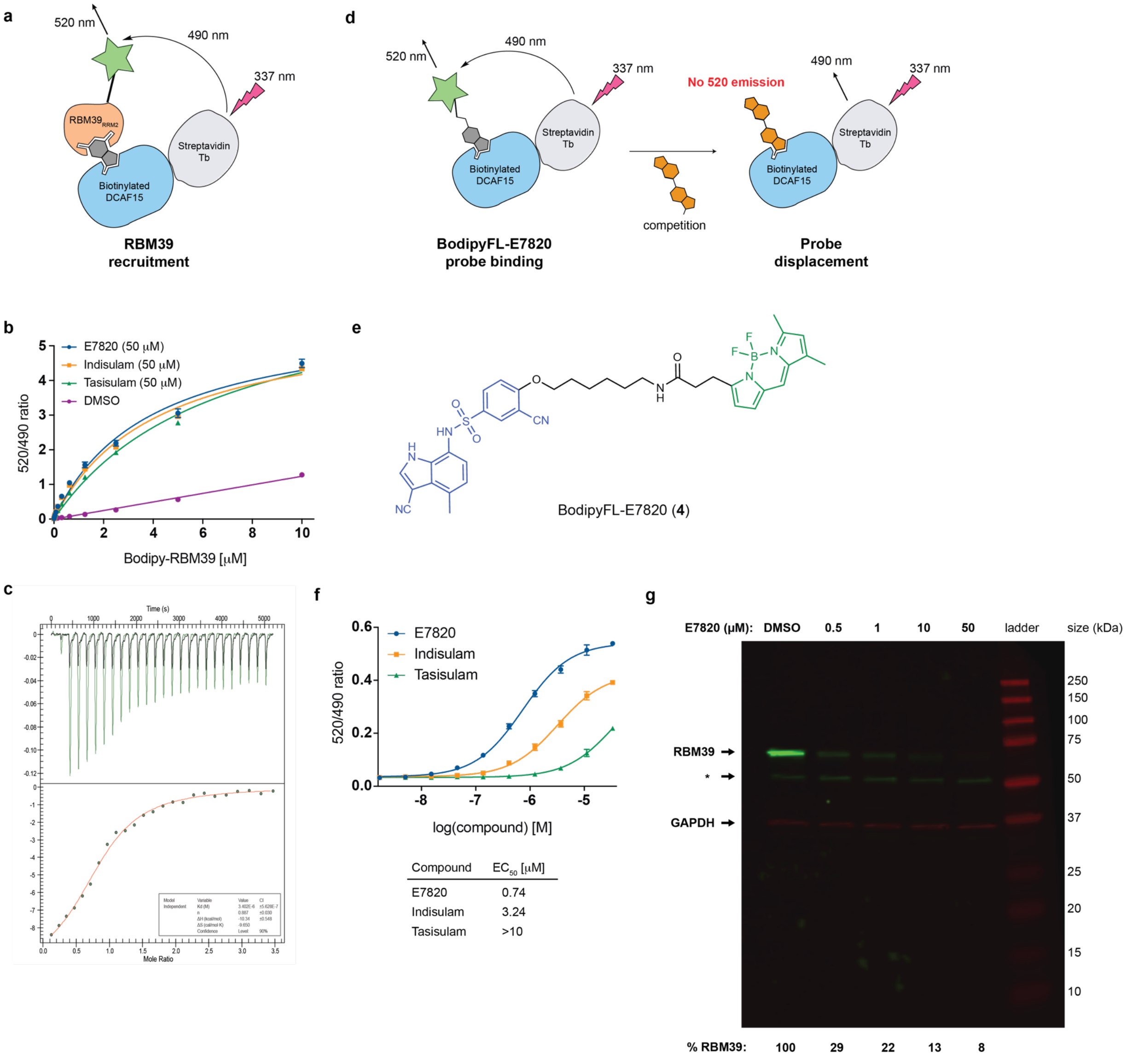


**Supplementary Figure 1. Biochemical characterization of DCAF15 binding to aryl-sulfonamides and RBM39$_{RRM2}$.** (**a**) Schematic representation of TR-FRET-based DCAF15-RBM39 dimerization assay. (**b**) TR-FRET. BodipyFL-RBM39$_{RRM2}$ titration (0.02-20 μM) to DDB1ΔB-DCAF15$_{biotin}$ at 200 nM in the absence (DMSO) or presence of aryl-sulfonamides at 50 μM. Data is plotted as means ± s.d. from three independent replicates (*n* = 3). (**c**) Assessment of E7820 binding to DDB1ΔB-DCAF15$_{biotin}$ using isothermal calorimetry. (**d**) Schematic representation of TR-FRET-based DCAF15 binding assay. Compound **4** is titrated to the DDB1ΔB-DCAF15$_{biotin}$, and the probe is displaced by competitor compounds. (**e**) Chemical structure of BodipyFL-E7820 (Compound **4**). (**f**) TR-FRET. Titration of E7820 ($EC_{50}$ = 0.74 µM), indisulam ($EC_{50}$ = 3.2 µM) or tasisulam ($EC_{50}$ > 10 µM) to DDB1ΔB-DCAF15$_{biotin}$ and BodipyFL-RBM39$_{RRM2}$. Data is plotted as means ± s.d. from three independent replicates (*n* = 3). (**g**) Cellular degradation of endogenous RBM39. HEK293T cells were treated with increasing concentrations of E7820 for 6 h, and protein levels were assessed by western blot with fluorescent secondary antibodies. Shown is the uncropped blot with molecular size markers. The positions of RBM39 and GAPDH are indicated, and the asterisk marks a non-specific band. Quantitation of RBM39 and GAPDH bands was performed with the Li-Cor imaging system, and %RBM39 is calculated as [RBM39 intensity / GAPDH intensity]. The experiment was performed once.

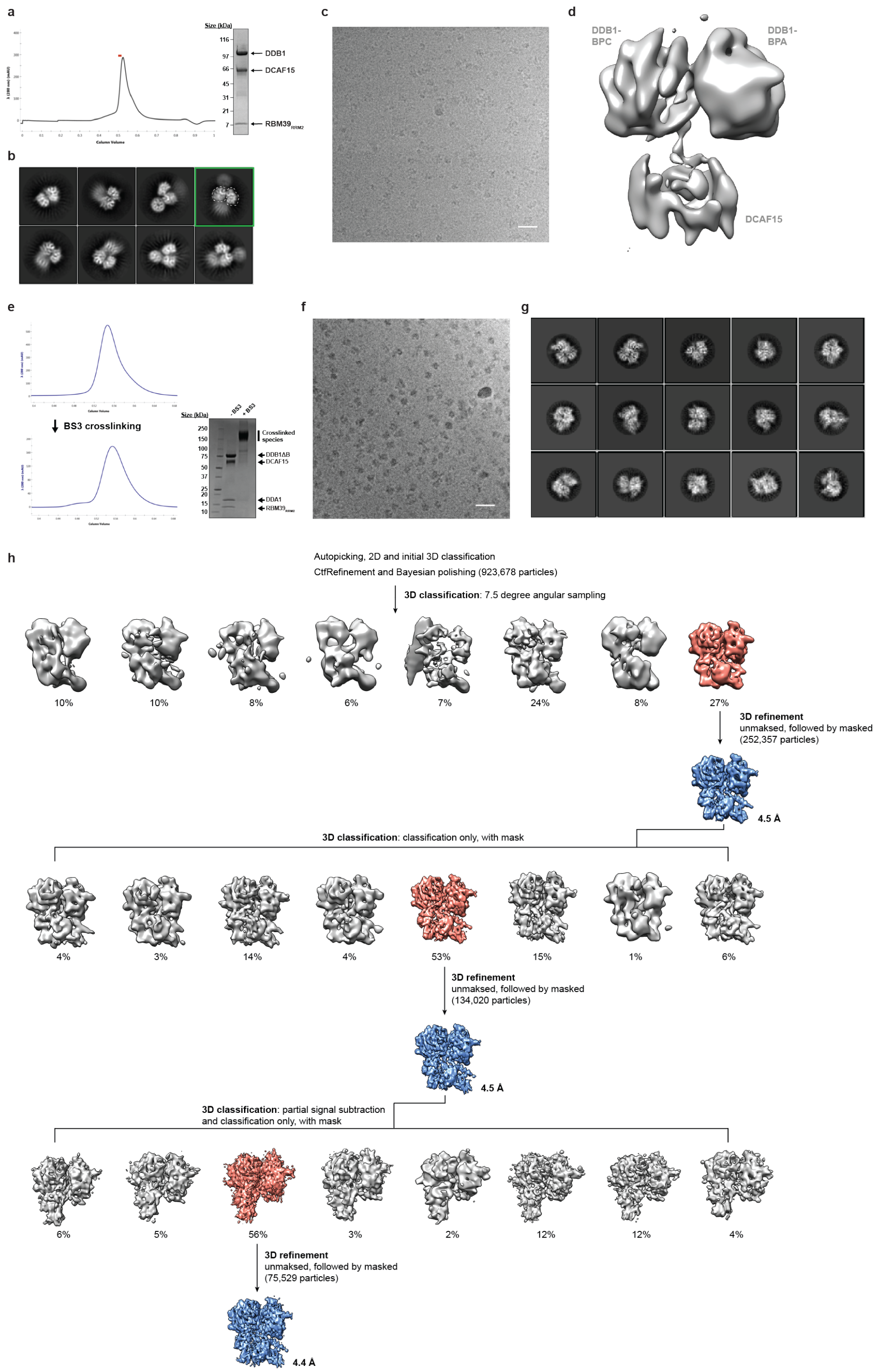


**Supplementary Figure 2. Cryo-EM of the 10 Å DDB1-DCAF15-RBM39$_{RRM2}$ complex bound to E7820 and the 4.4 Å DDB1ΔB-DCAF15-DDA1-RBM39$_{RRM2}$ complex bound to**

**E7820.** (**a**) Monodisperse peak of the complex by gel filtration. The red bar indicates the fraction used for the Coomassie-stained SDS-PAGE gel, shown to the right of the trace. Gel filtration of the complex was repeated at least three times with similar results. (**b**) Reference-free 2D class averages. The class average highlighted with a green box has signal for the three β-propellers of DDB1 and DCAF15, however only β-propellers A and C of DDB1 (outlined with a white hatched line) are aligned well, due to the inherent flexibility of DDB1 β-propeller B and DCAF15 (**c**) Representative cryo-EM micrograph at -2.3 μm defocus. Scale bar indicates 20 nm. Data collection was performed one time on this sample. (**d**) Reconstruction of the complex, highlighting the density for DDB1 β-propeller A, DDB1 β-propeller C, and DCAF15 at an average resolution of 10 Å. (**e**) The DDB1ΔB-DCAF15-DDA1-$RBM39_{RRM2}$ complex bound to E7820 complex displays a monodisperse gel filtration peak after BS3 cross-linking. The higher molecular weight cross-linked complex (~180 kDa) is indicated on the SDS-PAGE gel to the right. Cross-linking and gel filtration of this complex was performed at least three times with similar results. (**f**) Representative micrograph of the DDA1-containing complex imaged with a Volta phase plate (VPP) at -1.1 μm defocus. Scale bar represents 20 nm. Data collection was performed on this complex from two independent grids over the course of four imaging sessions. (**g**) Reference-free 2D class averages. (**h**) Data processing scheme for the crosslinked DDB1ΔB-DCAF15-DDA1-E7820-$RBM39_{RRM2}$ complex. Initial 2D and 3D classification resulted in 923,678 particles for Ctf Refinement and Bayesian polishing. Three subsequent rounds of 3D classification and refinement improved map resolution to a final average resolution of 4.4 Å. Percentages refer to the particles in each class. Red density maps indicate the classes that were used for the next round of processing, while blue density maps are from 3D refinements.

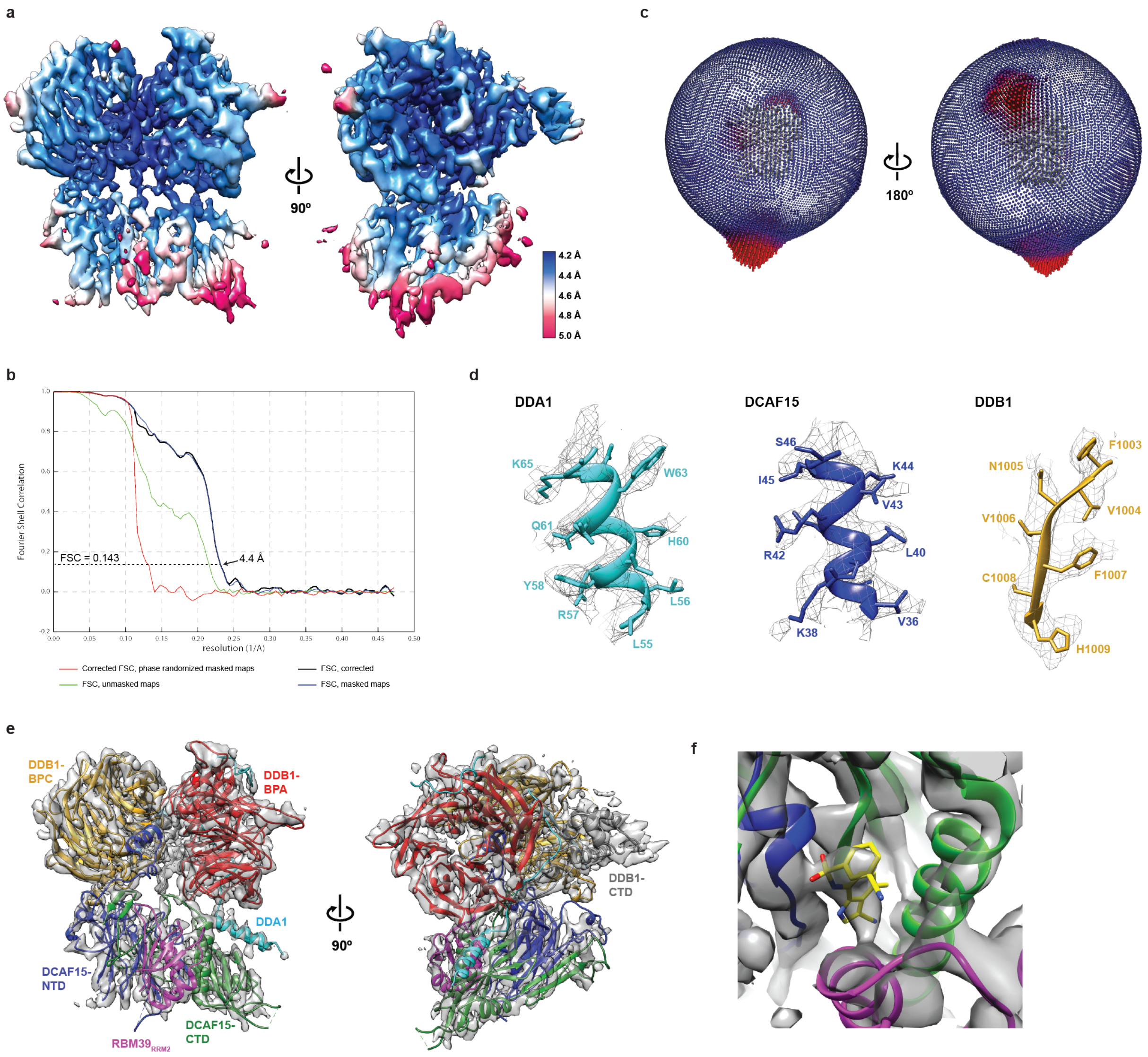


**Supplementary Figure 3. Local resolution, angular distribution, and model fitting of final cryo-EM reconstruction for DDB1ΔB-DCAF15-DDA1-E7820-RBM39$_{\mathrm{RRM2}}$.** (**a**) Local resolution map of final reconstruction calculated using Relion 3.0, colored according to the scale on the right. (**b**) Fourier shell correlation (FSC) plots for unmasked and masked maps, as well as phase randomized masked maps. Average resolution is indicated at FSC = 0.143. (**c**) Euler angle distribution of the 4.4 Å reconstruction in two views. (**d**) Regions of the cryo-EM model for DDA1, DCAF15, and DDB1 shown fit into the density from the sharpened map, demonstrating side chain density for multiple residues. Each density in mesh is shown at a threshold of 0.021 (from Chimera). (**e**) The crystal structure of DDB1ΔB-DCAF15$_{split}$-DDA1-RBM39$_{RRM2}$ bound to E7820 was docked and real space-refined into the unsharpened cryo-EM map of DDB1ΔB-DCAF15-DDA1-RBM39$_{RRM2}$ bound to E7820 using phenix dock in map and phenix realspace refine. Shown is the crystal structure (in cartoon) fit into the unsharpened cryo-EM density (gray surface). The majority of the cryo-EM density is accounted for by the crystal structure, both indicating that the cryo-EM map is also missing the flexible region between the DCAF15 NTD and CTD and that DCAF15$_{split}$ recapitulates the fold of full-length DCAF15. The cryo-EM density is shown at a threshold of 0.0145 (from Chimera). (**f**) Density for E7820 in the sharpened 4.4 Å cryo-EM reconstruction. Shown is the same docked model as in e, and the compound was placed by superimposing the E7820 bound crystal structure demonstrating density for E7820 (yellow) sandwiched between the DCAF15 NTD (blue), DCAF15 CTD (green), and RBM39$_{RRM2}$ (magenta). The sharpened cryo-EM density (B-factor of -129 from relion postprocessing) is shown in grey at a threshold of 0.0247 (from Chimera).

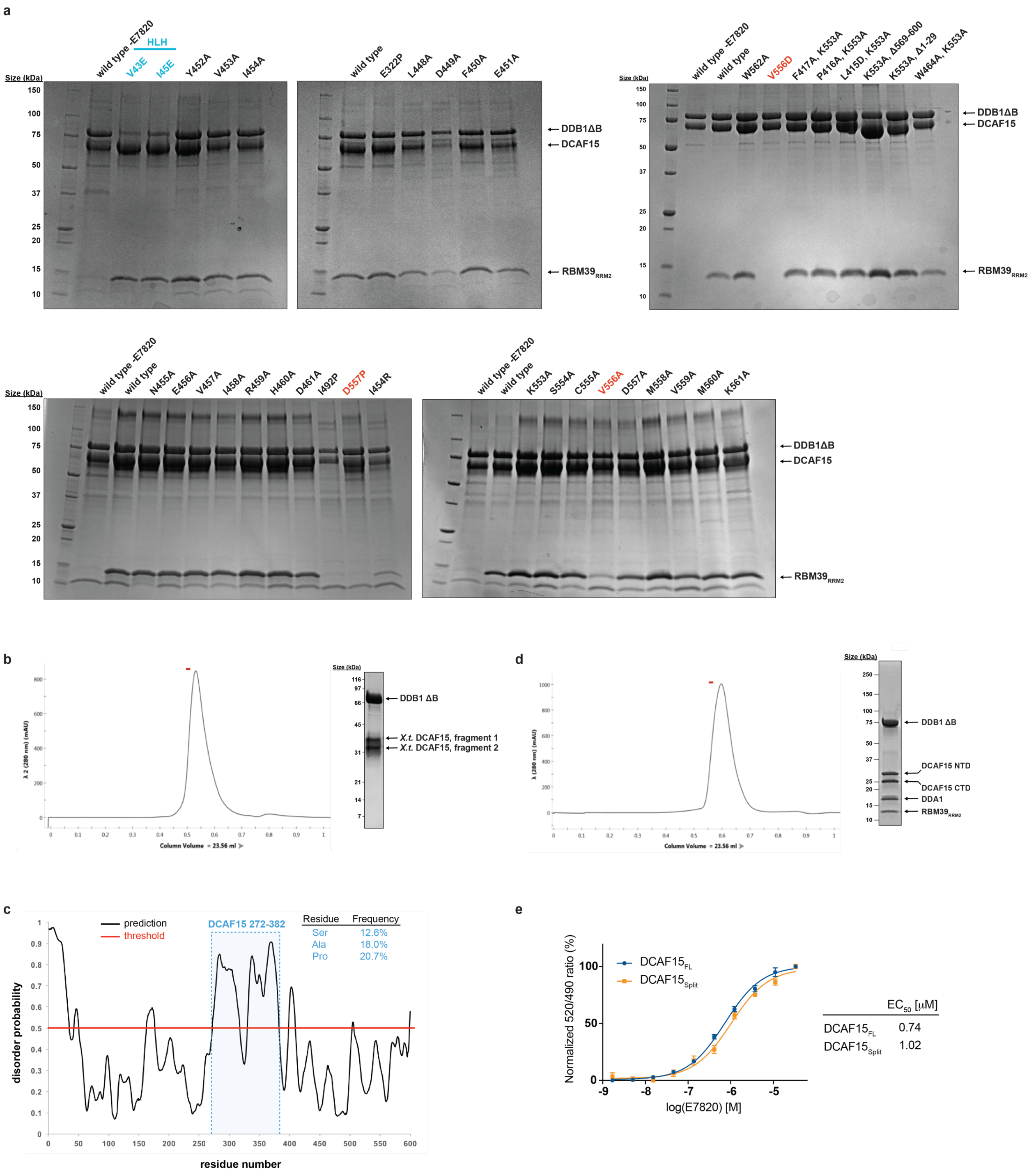


**Supplementary Figure 4. Mutant DCAF15 pull down and engineering a DCAF15$_{split}$ construct for crystallographic studies.** (**a**) Viruses expressing Strep II-tagged wild type and mutant DCAF15 were co-infected with viruses expressing DDB1ΔB and RBM39$_{RRM2}$ in Hi5

insect cells for 40 hours. STPEP purifications from Hi5 lysate were used to assess the interaction of DCAF15 mutants with DDB1ΔB and $RBM39_{RRM2}$. Unless indicated, infections and pull downs contained E7820. Mutants in the helix-loop-helix (HLH) domain are indicated with cyan text while mutants displaying an $RBM39_{RRM2}$ binding defect, without altering DCAF15 expression levels, are indicated in red. DCAF15 mutant pull downs were performed one time. (**b**) Limited proteolytic cleavage of the DDB1ΔB-*X.t.* DCAF15 complex with chymotrypsin followed by gel filtration demonstrates that two DCAF15 fragments, approximately 30 and 35 kDa in size, are associated with DDB1ΔB. Limited proteolysis on this complex was performed at least two times with similar results. (**c**) A plot of disorder prediction in DCAF15 using PrDOS with a 5% false positive rate. The plot indicates an internal region of DCAF15, residues 272-382, that is predicted to be disordered. The table inset to the right indicates that 50% of the internal region is composed of serine, alanine, and proline residues. (**d**) Representative gel filtration trace of the DDB1ΔB-$DCAF15_{split}$-DDA1-$RBM39_{RRM2}$ complex bound to E7820, demonstrating a monodispersed peak. The red bar indicates the fraction displayed on the gel to the right of the trace for both b and d. Gel filtration was performed at least three times with this complex with similar results. (**e**) TR-FRET. Titration of E7820 (0.002-33 μM) to full length DDB1ΔB-DCAF15 at 200 nM or DDB1ΔB-$DCAF15_{split}$ at 200 nM in the presence of BodipyFL-$RBM39_{RRM2}$ at 200 nM. DCAF15-RBM39 dimerization is measured and the data shows equivalent binding for full length DCAF15 and $DCAF15_{split}$. Data is plotted as means ± s.d. from three independent replicates ($n$ = 3).

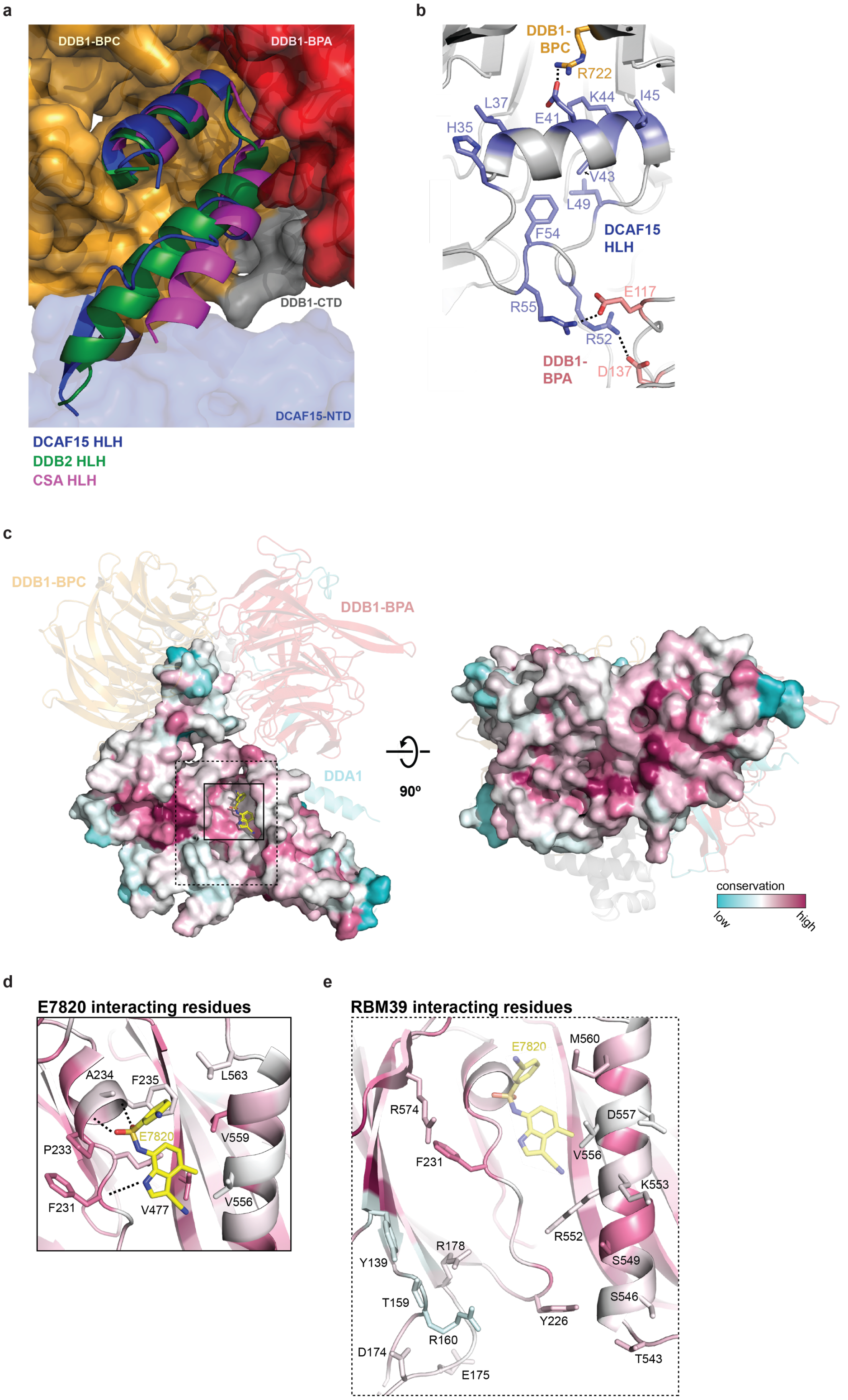

a
DDB1-BPC
DDB1-BPA
DDB1-CTD
DCAF15-NTD
DCAF15 HLH
DDB2 HLH
CSA HLH
b
DDB1-BPC
R722
L37
K44
I45
E41
H35
V43
L49
F54
DCAF15 HLH
E117
R55
R52
DDB1-BPA
D137
c
DDB1-BPC
DDB1-BPA
DDA1
90º
conservation
low
high
d
E7820 interacting residues
A234
F235
L563
E7820
V559
P233
F231
V477
V556
e
RBM39 interacting residues
E7820
M560
R574
D557
F231
V556
K553
R552
R178
S549
Y139
T159
S546
Y226
R160
D174
T543
E175

**Supplementary Figure 5. DCAF15 helix-loop-helix motif and conservation**. (**a**) Superposition of the CSA (pdb: 4a11) and DDB2 (pdb: 3ei4) helix-loop-helix (HLH) motif with DCAF15. The DDB1 BPA, BPC, and CTD are shown as a surface representation. (**b**) The DCAF15 HLH buries several hydrophobic residues, indicated in blue, between DDB1 BPA and BPC. Also shown are three salt bridges (black dotted lines) between DCAF15 E41, R55, and R52 and DDB1 R722, E117, and D137, respectively. (**c**) Overall conservation of DCAF15, shown in surface representation and colored according to the scale on the bottom right. DCAF15 conservation was analyzed by ConSurf[1]. DCAF15 sequences were first obtained with phmmer[2] using the full-length human DCAF15 sequence. An alignment of the sequences from phmmer was then used in hmmsearch to obtain more divergent DCAF15 orthologues. Finally, the 356 sequences from hmmsearch were aligned with Clustal Omega[3], and the multiple sequence alignment (MSA) was used in ConSurf. Shown in shaded cartoon representation are DDB1-BPA, DDB1-BPC, and DDA1. The black box outlines the E7820 interacting residues show in (**d**) and the dotted box outlines the RBM39 interacting residues shown in (**e**). (**d**) Conservation of the DCAF15 residues that interact with E7820. (**e**) Conservation of DCAF15 residues that interact with $RBM39_{RRM2}$.

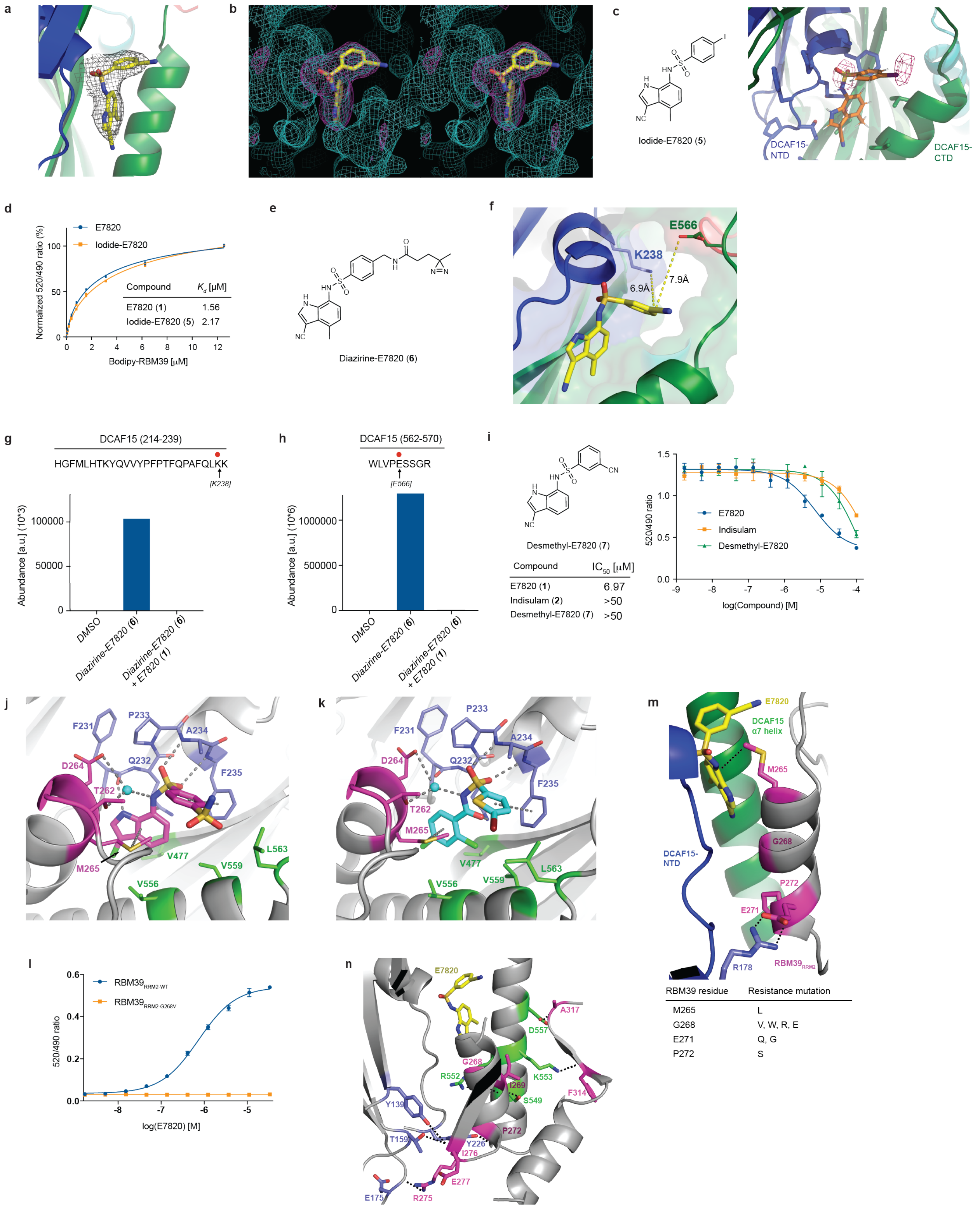
a
b
c
Iodide-E7820 (5)
DCAF15-NTD
DCAF15-CTD
d
E7820
Iodide-E7820
Normalized 520/490 ratio (%)
Bodipy-RBM39 [μM]
Compound
Kd [μM]
E7820 (1) 1.56
Iodide-E7820 (5) 2.17
e
Diazirine-E7820 (6)
f
E566
K238
6.9Å
7.9Å
g
DCAF15 (214-239)
HGFMLHTKYQVVYPFPTFQPAFQLKK
[K238]
Abundance [a.u] (10^3)
DMSO
Diazirine-E7820 (6)
Diazirine-E7820 (6) + E7820 (1)
h
DCAF15 (562-570)
WLVPESSGR
[E566]
Abundance [a.u] (10^6)
DMSO
Diazirine-E7820 (6)
Diazirine-E7820 (6) + E7820 (1)
i
Desmethyl-E7820 (7)
Compound IC50 [μM]
E7820 (1) 6.97
Indisulam (2) >50
Desmethyl-E7820 (7) >50
520/490 ratio
E7820
Indisulam
Desmethyl-E7820
log(Compound) [M]
j
F231
P233
A234
Q232
F235
D264
T262
M265
V477
V556
V559
L563
k
F231
P233
A234
D264
Q232
F235
T262
M265
V477
V556
V559
L563
m
E7820
DCAF15 α7 helix
M265
G268
P272
E271
DCAF15-NTD
R178
RBM39 RRM2
RBM39 residue | Resistance mutation
M265 | L
G268 | V, W, R, E
E271 | Q, G
P272 | S
l
RBM39 RRM2-WT
RBM39 RRM2-G268V
520/490 ratio
log(E7820) [M]
n
E7820
A317
D557
G268
R552
K553
I269
S549
F314
Y139
P272
T159
Y226
I276
E277
E175
R275

**Supplementary Figure 6. Experimental validation of E7820 binding site and resistance mutations in RBM39. (a)** Shown in mesh is the 2Fo-Fc electron density map for E7820, contoured at 1.0 sigma **(b)** Stereo view of a simulated annealing omit map around E7820 in magenta and 2Fo-Fc in cyan. (**c**) Crystal structure of DDB1ΔB-DCAF15$_{split}$-DDA1-RBM39$_{RRM2}$ bound to compound **5** (Iodide-E7820). Shown in mesh is the anomalous difference map contoured at 5 sigma. (**d**) TR-FRET. Titration of BodipyFL-RBM39$_{RRM2}$ to DDB1ΔB-DCAF15$_{biotin}$ at 200 nM pre-treated with E7820 or compound **5** at 50 μM ($n$ = 2). (**e**) Chemical structure of compound **6** (Diazirine-E7820), used for UV-crosslinking. (**f**) Two UV-crosslinked residues (Lys238, Glu566) in DCAF15 are highlighted. The distances from the 4-position of E7820 phenyl ring to the residues are specified (6.9 Å and 7.9 Å, respectively). (**g-h**) UV-crosslinking coupled with mass spectrometry. Proteins were treated by DMSO, compound **6**, or compound **6** with pre-treatment of E7820 then UV-irradiated (see **Online Methods**). (**g**) Quantification of modified peptides in DCAF15 214-239. Red circle above the sequence indicates the UV-crosslinked residue (Lys238). (**h**) Quantification of modified peptides in DCAF15 562-570. Red circle above the sequence indicates the UV-crosslinked residue (Glu566). (**i**) TR-FRET based competitive binding assay of compound **7** (Desmethyl-E7820). Data is plotted as means ± s.d. from three independent replicates ($n$ = 3). (**j**) Side chain interactions between DCAF15 NTD (blue), DCAF15 CTD (green), RBM39$_{RRM2}$ (magenta), water (cyan sphere), and indisulam (magenta compound). (**k**) Side chain interactions between DCAF15 NTD, DCAF15 CTD, RBM39$_{RRM2}$, water, and tasisulam (cyan), using the same protein coloring as (**j**). (**l**) TR-FRET. Titration of E7820 (0.002-33 μM) to full length DDB1ΔB-DCAF15 at 200 nM in the presence of BodipyFL-RBM39$_{RRM2-WT}$ ($EC_{50}$ = 0.74 μM) or BodipyFL-RBM39$_{RRM2-G268V}$ at 200 nM. Data is plotted as means ± s.d. from three independent replicates ($n$ = 3) (**m**) RBM39 resistance mutations. Labeled in magenta are the four positions in RBM39 that confer resistance to indisulam-dependent toxicity when mutated[4]. The table below indicates the resistance mutations at these positions. (**n**) Shown are the network of residues involved in backbone hydrogen bonds between RBM39 (magenta) and DCAF15 (NTD residues in blue and CTD residues in green).

### References


1 Landau, M. *et al.* ConSurf 2005: the projection of evolutionary conservation scores of residues on protein structures. *Nucleic acids research* **33**, W299-302, doi:10.1093/nar/gki370 (2005).

2 Potter, S. C. *et al.* HMMER web server: 2018 update. *Nucleic Acids Res* **46**, W200-W204, doi:10.1093/nar/gky448 (2018).

3 Larkin, M. A. *et al.* Clustal W and Clustal X version 2.0. *Bioinformatics (Oxford, England)* **23**, 2947-2948, doi:10.1093/bioinformatics/btm404 (2007).

4 Han, T. *et al.* Anticancer sulfonamides target splicing by inducing RBM39 degradation via recruitment to DCAF15. *Science* **356**, doi:10.1126/science.aal3755 (2017).
